# Lrrcc1 and Ccdc61 are conserved effectors of multiciliated cell function

**DOI:** 10.1101/2021.01.30.428946

**Authors:** Aude Nommick, Camille Boutin, Olivier Rosnet, Elsa Bazellières, Virginie Thomé, Etienne Loiseau, Annie Viallat, Laurent Kodjabachian

## Abstract

Ciliated epithelia perform a variety of essential functions across animal evolution, ranging from locomotion of marine organisms to mucociliary clearance of airways in mammals. These epithelia are composed of multiciliated cells (MCCs) harbouring myriads of motile cilia, which rest on modified centrioles called basal bodies (BBs), and beat coordinately to generate directed fluid flows. Thus, BB biogenesis and organization is central to MCC function. In basal eukaryotes, the coiled-coil domain proteins Lrrcc1 and Ccdc61 were shown to be required for proper BB construction and function. Here, we used the *Xenopus* embryonic ciliated epidermis to characterize Lrrcc1 and Ccdc61 in vertebrate MCCs. We found that they both encode BB components, with a prominent association to striated rootlets. Knocking down either gene caused defects in BB docking, spacing, and polarization. Moreover, their depletion impaired the apical cytoskeleton, and altered ciliary beating. Consequently, cilia-powered fluid flow was greatly reduced in morphant tadpoles, which displayed enhanced mortality when exposed to pathogenic bacteria. This work illustrates how integration across organizational scales make elementary BB components essential for the emergence of the physiological function of ciliated epithelia.

## Introduction

Multiciliated epithelia are composed of multiciliated cells (MCCs) harboring numerous motile cilia. Ciliary beating generates powerful strokes essential for a variety of physiological functions in animals (Meunier and Azimzadeh, 2016). In aquatic organisms of the *Lophotrochozoan* and *Echinodermata* phyla, coordinated MCC beating is required for locomotion, clearance and transport of particles, and for feeding of larvae. In vertebrates, MCCs produce external or internal fluid flows. In lungfish, the ciliated epidermis clears the animal of particles and settling organisms before hatching (Kemp, 1996). In amphibian embryos, several roles have been proposed for the ciliated epidermis: prevention of micro-organisms and debris from attaching to the epidermis, pre-hatching rotation and post-hatching gliding, respiratory gas exchange, movement of surface mucus films and transportation of chemical signals to the olfactory organs (Nokhbatolfoghahai et al., 2006). Which of those roles is carried out by MCCs of the *Xenopus* embryonic epidermis remains unclear, despite a recent wealth of mechanistic studies in this model (Boutin and Kodjabachian, 2019). In mammals, among other functions, MCCs help circulation of the cerebrospinal fluid in the central nervous system, mucociliary clearance of pathogens and pollutants from airways, and transportation of gametes in genital tracts (Spassky and Meunier, 2017). Consequently, mutations in genes necessary for multiple cilia formation or beating cause familial syndromes characterized by severe chronic airway infections, and an elevated risk of infertility (Spassky and Meunier, 2017). A precise multiscale organization of ciliary beating is required to establish a robust and directed flow at the surface of ciliated epithelia. At the cellular scale, all cilia must beat in the same direction, and at the tissue scale the beating direction must be coordinated between neighboring cells. The Planar Cell Polarity pathway plays a prominent role in the establishment and maintenance of cilia polarity, both at tissue and cellular scales (Boutin et al., 2014; Chien et al., 2015; Mitchell et al., 2009; Park et al., 2008; Walentek et al., 2017).

Motile cilia of MCCs are built via a multistep process (Boutin and Kodjabachian, 2019; Spassky and Meunier, 2017). First, numerous centrioles must be produced and subsequently released in the cytoplasm. Next, centrioles migrate and dock at the apical surface, where they acquire a regular distribution and a coordinated orientation. Finally, ciliary axonemes extend from docked centrioles, and metachronal waves of ciliary beating are initiated and subsequently reinforced by mechanical feedback from the flow, which refines the coordination of cilia polarity. During their journey towards the surface, centrioles mature into basal bodies (BBs) by acquiring basal foot (BF) and rootlet appendages, which localize asymmetrically and are essential for cilia polarization (Meunier and Azimzadeh, 2016). In *Xenopus* epidermal MCCs, two different types of rootlets attach to the proximal end of BBs. The most prominent rootlet has a fan shape and is localized opposite to the BF, which itself is positioned distally on the BB. The second rootlet is much longer and thinner and dives into the cytoplasm (Zhang and Mitchell, 2015). These appendages confer an intrinsic polarity to BBs, which in mature MCCs reflects the direction of ciliary beating, with the BF pointing in the direction of the effective stroke.

MCC cilia formation and organization relies on close interactions between BBs and cytoskeletal elements (Boutin and Kodjabachian, 2019; Meunier and Azimzadeh, 2016). The transport and docking of centrioles/BBs to the apical surface is dependent on acto-myosin-based mechanisms (Boisvieux-Ulrich et al., 1990; Epting et al., 2015; Kulkarni et al., 2018; Lemullois et al., 1988; Miyatake et al., 2015; Park et al., 2008). Once docked, neighboring BBs are linked by subapical actin filaments and apical microtubules (MTs) emanating from rootlets and BFs, respectively. The geometrical network hence made ensures regular BB spacing and coordinated orientation over the apical cell surface (Antoniades et al., 2014; Lemullois et al., 1988; Park et al., 2006; Werner et al., 2011; Yasunaga et al., 2015). Chemical interference with actin or MT networks leads to BB disorganization and impaired ciliary function (Herawati et al., 2016; Werner et al., 2011). Reciprocally, depletion of specific centriolar components, which prevents the formation of appendages and/or preclude BB/cytoskeleton interactions, alter BB organization and ciliary function (Antoniades et al., 2014; Bustamante-Marin et al., 2019; Clare et al., 2014; Herawati et al., 2016; Kulkarni et al., 2018; Kunimoto et al., 2012; Turk et al., 2015; Walentek et al., 2016).

Lrrcc1 (Leucine rich repeat coiled-coil domain containing 1) and Ccdc61 (Coiled-coil domain-containing protein 61) are structural proteins, conserved from *Chlamydomonas* to human, which are involved in centriole appendage biogenesis and function, both at the centrosome and at ciliary BBs (Adams et al., 1985; Barenz et al., 2018; Basquin et al., 2019; Bengueddach et al., 2017; Hoops et al., 1984; Muto et al., 2008; Ochi et al., 2020; Pizon et al., 2020; Silflow et al., 2001; Wright et al., 1983). In MCCs of the planarian epidermis, Lrrcc1 (Vfl1) and Ccdc61 (Vfl3) depletion causes structural BB defects, thus perturbing cilia orientation, and altering the direction of locomotion (Basquin et al., 2019). The importance of Lrrcc1 and Ccdc61 in vertebrate MCCs remains unknown. Here, we used the *Xenopus laevis* ciliated epidermis as a model to shed light on this issue. We decided to comparatively study Lrrcc1 and Ccdc61, based on their shared biological functions in *Chlamydomonas* and *Schmidtea* (Adams et al., 1985; Basquin et al., 2019; Hoops et al., 1984; Wright et al., 1983), and their reported physical interaction in an unbiased human proteomic screen (Hein et al., 2015). In *Xenopus*, MCCs are specified deeply, in the inner cellular layer of the epidermis before intercalating at regular intervals into the outer cellular layer. BB synthesis is initiated while MCCs are still in the inner epidermal layer; BB docking, distribution, orientation and ciliogenesis are completed when MCCs have radially intercalated and expanded their apical surface.

We report here that Lrrcc1 and Ccdc61 are both associated to *Xenopus* MCC centrioles, with a preferential localization at rootlets of mature BBs. We found that knocking down either gene impacts ciliated epithelium biogenesis at multiple scales. At the organelle scale, Ccdc61 is required for rootlet association of Pericentrin (Pcnt), that we characterize as a novel marker of this appendage in *Xenopus* BBs. At the cellular scale, we show that Lrrcc1 and Ccdc61 are required for proper organization of BBs. In addition, MCCs depleted for either gene present important defects in apical cytoskeleton organization. Finally, at the scale of the embryo, Lrrcc1 and Ccdc61 knock-down revealed their importance for the generation of a directed fluid flow and the resistance to opportunistic pathogens. This study bridges multiple scales of analysis to reveal how intracellular disorganization of MCCs can impair the physiology of the whole organism.

## Results

### Lrrcr1 and Ccdc61 are associated to centrioles and basal bodies in MCCs

Single cell RNA sequencing (scRNA-seq) of *Xenopus tropicalis* embryos revealed that *lrrcc1* and *ccdc61* are specifically expressed in ciliated epidermal cells from gastrula stages onwards (Briggs et al., 2018), and both genes were found to be activated by the Multicilin/E2F4/5 complex, which is known to be necessary and sufficient for vertebrate MCC differentiation (Ma et al., 2014). We used whole mount *in situ* hybridization to analyze the localization of *lrrcc1* and *ccdc61* transcripts in *Xenopus laevis* embryos. At early tailbud stage 20, both genes displayed a “salt and pepper” pattern typical of epidermal MCCs (Fig. S1A). Next, we analyzed protein subcellular localization at different time points during MCC differentiation. First, we used antibody staining to analyze the distribution of the endogenous Lrrcc1 protein. At stage 16, Lrrcc1 was detected in the vicinity of Centrin-positive cytoplasmic clouds, where centriole synthesis occurs (Fig. 1A). At stage 18, when released centrioles start migrating towards the apical surface, Lrrcc1 appeared closely associated to individual neo-centrioles marked by Centrin (Fig. 1B). At stage 31, Lrrcc1 was associated to mature BBs docked at the apical surface. Interestingly, the Lrrcc1 signal was systematically located on one side of Centrin-positive BBs. In orthogonal view, the signal flanked the BB and extended towards the cytoplasm (Fig. 1C). This localization suggested that Lrrcc1 associates to the BB rootlet. To further confirm this, we carefully analyzed the 3D distribution of Lrrcc1 relative to Centrin and γ-Tubulin, known to localize to the BF cap in MCCs (Clare et al., 2014; Hagiwara et al., 2000). On a top view, Centrin appeared as a single dot (Fig. 1D and Fig. S1B). Two pools of γ-Tubulin were observed juxtaposed to the BB. The first one appeared as a dot with strong intensity juxtaposed to the BB core. A lateral view revealed that this dot was located at a depth similar to that of the BB (Fig. 1D1, Fig. S1B). This pool corresponds to the γ-Tubulin previously described at the BF. In contrast, the second pool of γ-Tubulin displayed a less intense signal. It appeared as one or two dots localized opposite to the BF with respect to the BB (Fig. S1B). This pool localized deeper, extending from the BB into the cytoplasm, in a position compatible with the double rootlets described in *Xenopus* MCCs (Zhang and Mitchell, 2015). The Lrrcc1 signal appeared either as a single dot (Fig. 1D1), or as double dots (Fig. 1D2), below the Centrin signal and extending towards the cytoplasm. This analysis revealed that the endogenous Lrrcc1 protein localized to the two types of rootlets of *Xenopus* epidermal MCCs. It also revealed the association of γ-Tubulin to these appendages.

**Figure 1.**
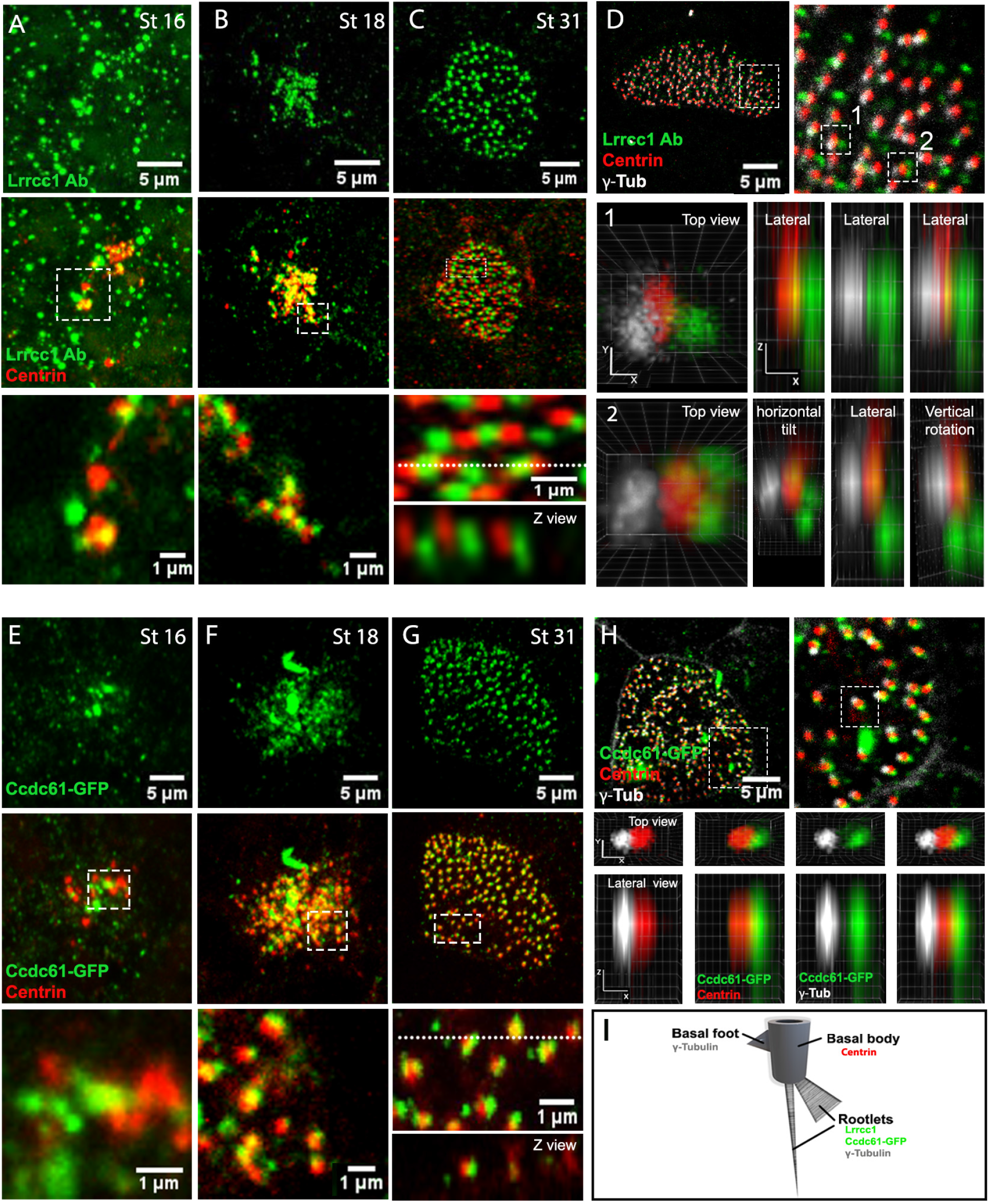
Llrcc1 and Ccdc61 associate with centrioles in multiciliated cells. (A-C) MIPs (maximum intensity projections) of confocal acquisitions of MCCs stained with Lrrcc1 (green) and Centrin (red) antibodies. White dashed boxes indicate higher magnification views presented below and the dashed white line sets the position of the Z view. (D) MIPs of confocal acquisitions showing the localization of Lrrcc1 (green) compared to Centrin (red) and γ-Tubulin (white). White dotted boxes 1 and 2 indicate the BBs analysed in 3D with Clear Volume. (E-G) MIPs of confocal acquisitions from MCCs expressing Ccdc61-GFP (green) and Centrin-RFP (red) fusion proteins. (H) Clear volume 3D top and lateral views of a BB labelled with Ccdc61-GFP and Centrin-RFP. (I) Scheme of a mature BB in a *Xenopus* epidermal MCC recapitulating the localization of proteins analysed in this figure.

The lack of cross-reacting antibodies prevented us from analyzing the endogenous Ccdc61 protein in *Xenopus* MCCs. Therefore, we injected a synthetic transcript coding for a Ccdc61-GFP fusion protein to get an insight on the putative distribution of endogenous Ccdc61 during MCC differentiation. At all stages analyzed, Ccdc61-GFP was observed in close vicinity of centrioles (Fig. 1E-G). In mature MCCs, triple staining with Centrin and γ-Tubulin indicated that Ccdc61-GFP was found opposite to the BF, in a position compatible with the proximal part of the rootlet (Fig. 1H). A similar localization was recently reported for a Ccdc61-RFP fusion in *Xenopus* epidermal MCCs (Ochi et al., 2020). We conclude that Lrrcc1 and Ccdc61 preferentially associate to rootlets in mature epidermal MCCs of *Xenopus*.

Altogether these data suggest that Lrrcc1 and Ccdc61 are associated to epidermal MCC centrioles from the time of their release in the cytoplasm through the phases of docking, BB maturation, ciliary growth and maintenance.

### Lrrcc1 and Ccdc61 depletion impairs centriole docking, spacing and orientation

The subcellular distribution of Lrrcc1 and Ccdc61 during MCC differentiation suggested that they could be involved in BB apical docking as well as distribution and orientation at the apical surface. To investigate this, we knocked down *lrrcc1* and *ccdc61* through injection in the presumptive epidermis of two independent morpholino antisense oligonucleotides (MOs) designed to block translation (MO-ATG) or splicing (MO-Spl) (Fig. S2A). The efficiency of MOs was established through the reduction of immunofluorescent signal intensity of endogenous Lrrcc1, or co-injected Ccdc61-GFP (Fig. S2B, C). We performed immunostaining at stage 31 to analyze BB organization. As previously described (Werner et al., 2011), BBs displayed a stereotypical organization in mature control MCCs. They were all docked, quite evenly distributed at the apical surface and oriented in a coordinated manner (Fig. 2A, C, E). In contrast, Lrrcc1 and Ccdc61 knock-down drastically impaired centriole organization and morphant cells could be classified in two phenotypic categories. The first phenotypic class was characterized by clusters of centrioles stuck in the upper half of the cytoplasm (Fig. 2A, Fig. S3A). To quantify this defect, we analyzed the apico-basal (A-B) distribution of centrioles along the Z-axis of confocal stack acquisitions. In control cells, most centrioles localized within the first 1,6 μm below the apical cell surface (Fig. 2B). In contrast, in Lrrcc1-ATG and Ccdc61-ATG morphant cells most centrioles localized deeper, between 1,6 μm and 5.6 μm from the surface (Fig. 2B). Similar results were obtained with Lrrcc1-Spl and Ccdc61-Spl MOs (Fig. S3B). In the second phenotypic class, most centrioles properly localized at the apical surface, but displayed irregular spacing and a randomized orientation (Fig. 2C, E and Fig. S3C, E). To quantify BB spacing defects, we applied Delaunay triangulation between the centroids of docked centrioles and measured the area of the obtained triangles. In the control situation, the even distribution of BBs resulted in many triangles of similar area that tightly distributed around the median (Fig. 2C, D and Fig. S3C, D). In contrast, in morphant cells, variable distances between BBs resulted in a broader distribution of triangles around the median (Fig. 2C, D and Fig. S3C, D). We revealed BB orientation by immunostaining the BB core (Centrin) and the offset BF (intense γ-Tubulin spot), which allowed us to automatically extract orientation vectors and plot their circular distribution. In control cells, vector angles tightly distributed around the mean, indicating that BBs oriented in the same direction. In Lrrcc1 and Ccdc61 morphant cells, the intense γ-Tubulin signal was largely preserved, suggesting that the integrity of the BF was maintained. However, vector angles were widely distributed around the mean, indicating a randomization of BB orientation (Fig. 2E and Fig. S3E). Accordingly, circular standard deviation values were significantly higher in morphants, as compared to control (Fig. 2F and Fig. S3F). The Rayleigh statistical test also revealed a higher percentage of morphant MCCs, in which no significant mean vector could be defined, as compared to control (Fig. 2G and Fig. S3G).

**Figure 2.**
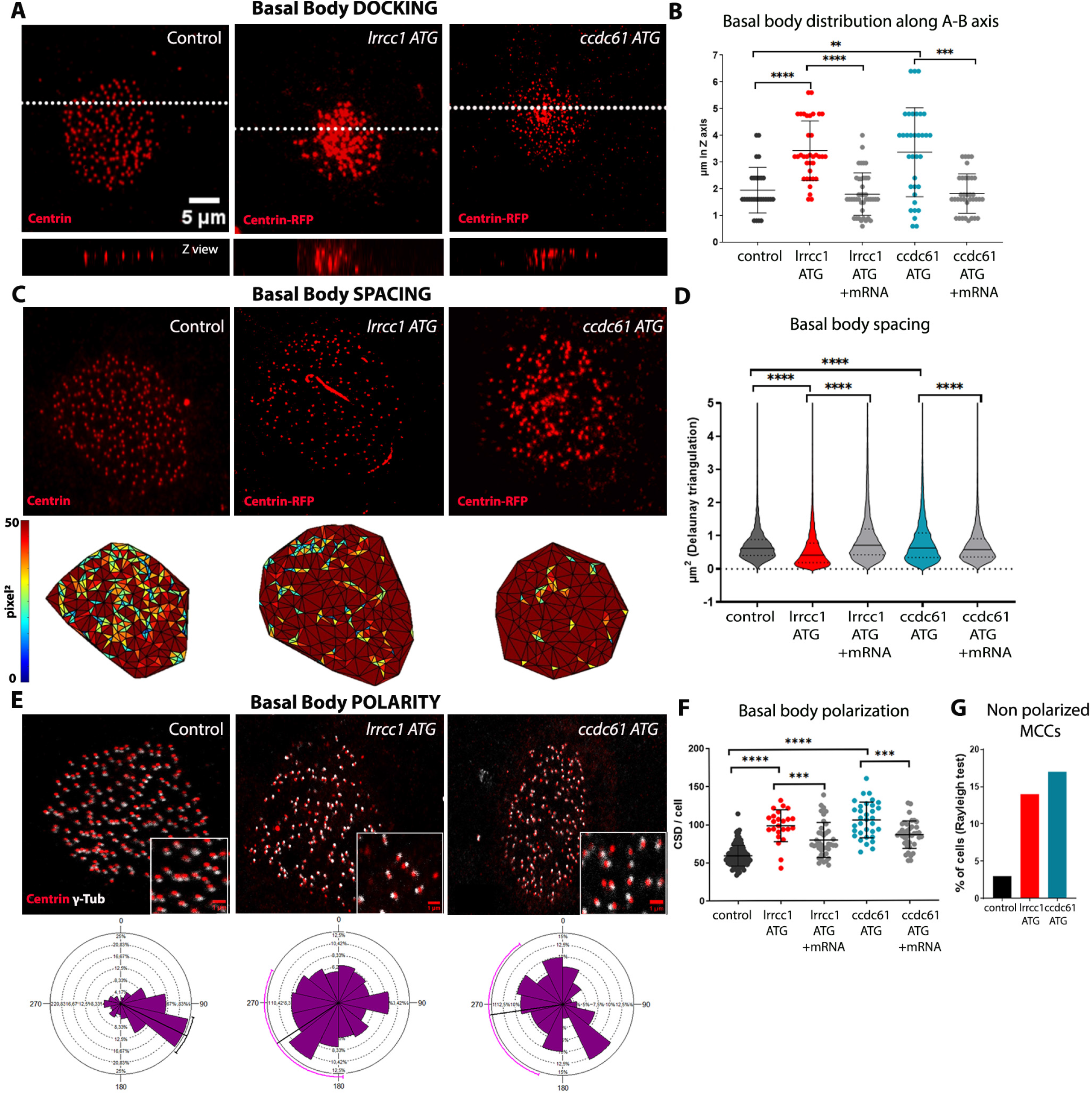
Lrrcc1 and Ccdc61 are necessary for BB docking, spacing and orientation. (A) MIPs of confocal acquisitions of MCCs from St 31 control, Lrrcc1 or Ccdc61 morphant embryos stained with Centrin Ab, or expressing Centrin-RFP, which was co-injected with MOs. (B) Graph displaying the apico basal (A-B) distribution of centrioles within control, Lrrcc1 and Ccdc61 morphant and rescued MCCs. Each point represents a single cell. Horizontal lines represent the mean and SD. (C) MIPs of confocal acquisitions of apical BBs in MCCs from St 31 control, Lrrcc1 or Ccdc61 morphant embryos stained with Centrin Ab, or expressing Centrin-RFP, which was co-injected with MOs. The corresponding Delaunay triangulation outputs are presented on the bottom row. Each colour is associated to a range of triangle areas in pixel^2^. (D) Violin plots displaying the distribution of triangle areas in μm^2^ of control, morphant and rescued MCCs. The horizontal line indicates the median, dashed lines indicate the quartiles. (E) MIPs of confocal acquisitions of control and morphant MCCs stained with Centrin (red, centrioles) and γ-Tubulin (white, BF). Below each cell, its respective rose histogram representing the distribution of BB orientations. The black line running from the centre of the diagram to the outer edge is the mean angle and the arcs extending to either side represent the confidence limits of the mean fixed at 95% (when the mean angle is only theoretical but not significant, the line turns pink). (F) Circular Standard Deviations of control, morphant and rescued MCCs. Each point represents a single cell. Horizontal lines represent the mean and SD. (G) Graph displaying the percentage of non-polarized MCCs (no significant mean angle of BBs within a cell can be calculated) following the Rayleigh statistical test. All confocal images are at the scale shown in (A).

In both Lrrcc1 and Ccdc61 morphant MCCs, cilia could be observed suggesting that those proteins are not required for ciliary growth *per se* (Fig. S4). Importantly, for both genes and both types of MOs, docking, spacing and orientation phenotypes were rescued by co-injection of *lrrcc1* or *ccdc61* mRNA constructs lacking (*lrrcc1*), or silently mutated (*ccdc61*) on the MO-binding sequence (Fig. 2B, D, F, Fig. S3B, D, F and S5).

Altogether, these data show that Lrrcc1 and Ccdc61 are required for correct apical migration and/or docking of BBs, as well as for their proper planar distribution and orientation at the apical surface.

### Ccdc61 is required for Pericentrin association to rootlet

Our localization and functional data suggested that Lrrcc1 and Ccdc61 could participate to the formation of rootlet appendages in *Xenopus* MCCs. To analyze this possibility, we performed TEM experiments on stage 31 MCCs from control and morphant embryos. The presence of centriole docking defects was used as a way to ascertain the morphant character of scored MCCs (Fig. 3B). Both fan-shape and thin rootlets could be observed in similar proportions in Lrrcc1 and Ccdc61 morphant cells, as compared to control (Fig. 3A, C). Whenever visible, rootlets did not present obvious structural or positioning defects.

**Figure 3.**
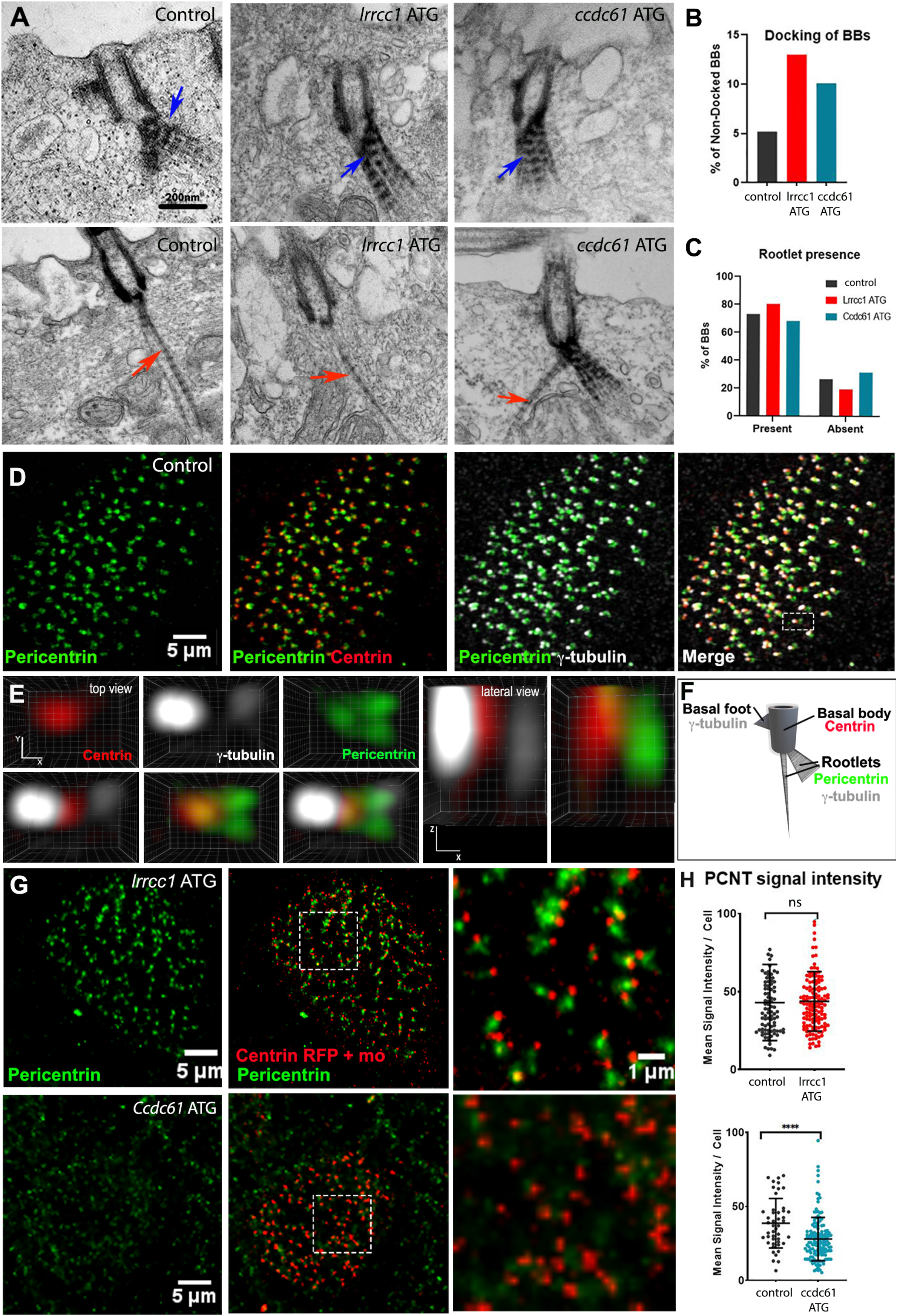
Ccdc61 is required for Pericentrin association to rootlet appendages. (A) Transversal TEM acquisitions of BBs from St 31 control and morphant MCCs. Both fan-shape (blue arrow) and long (red arrow) rootlets could be observed in all conditions, but rarely on the same section. (B) Quantification of BB docking on TEM acquisitions to corroborate morpholino efficiency. (C) Graph displaying the proportion of BBs with at least one (fan-shaped or long) or without rootlets quantified on TEM acquisitions. Please note that about 20% control BBs appear to lack rootlets due to the angle of the section. The same proportion was observed in Lrrcc1 and Ccdc61 morphant MCCs. (D) MIPs of confocal acquisitions of MCCs stained with Pcnt (green), Centrin (red) and γ-Tubulin (white). The white dashed box indicates the BB analysed in 3D in E. (E) Clear Volume 3D top and lateral views of a single BB. (F) Scheme of a mature BB recapitulating the localization of proteins analysed in this figure. (G) MIPs of confocal acquisitions of St 31 Lrrcc1 and Ccdc61 morphant MCCs expressing Centrin-RFP (red) and stained with Pcnt antibody (green). White dashed boxes indicate high magnification views displayed on the right. (H) Quantification of Pcnt mean signal intensity. Each point represents an MCC. Horizontal lines represent the mean and SD.

To look for potential molecular defects, we searched for an endogenous rootlet marker to be used in fluorescent imaging. Using home-made antibodies (Fig. S6A, B), we identified *Xenopus* Pcnt as a *bona fide* rootlet component. In stage 31 control cells, Pcnt immunostaining presented a dotted pattern, and was associated with Centrin and γ-Tubulin at the apical surface (Fig. 3D). Analysis of the relative distribution of these proteins in individual BBs revealed that Pcnt was located opposite to the strong BF-associated dot of γ-Tubulin, in a plane below Centrin. It was present as one or two dots emerging from the BB and extending towards the cytoplasm (Fig 3E, S6C). This analysis revealed that in *Xenopus* epidermal MCCs, Pcnt specifically localizes at rootlets.

Next, we analyzed Pcnt signal in Lrrcc1 and Ccd61 morphant embryos. To avoid immunostaining variability between different embryos, we compared Pcnt signal intensity within mosaic embryos. For Lrrcc1, no differences were observed between non-injected and morphant cells, which, however, clearly displayed randomized BB polarity (Fig. 3G-H). In contrast, a marked decrease of Pcnt signal was observed in Ccdc61 morphant cells, as compared to non-injected cells from the same embryos (Fig. 3G-H).

Altogether, these experiments suggest that Lrrcc1 and Ccdc61 are not required for building rootlet appendages in *Xenopus* MCCs. However, Ccdc61 is required for association of the protein Pcnt to the rootlet, which in turn could affect rootlet function and BB organization.

### Lrrcc1 and Ccdc61 depletion disturbs apical cytoskeleton organization

BB organizational defects observed in Ccdc61 and Lrrcc1 morphant cells could be linked to defective apical cytoskeleton. To test this idea, we first analyzed F-actin networks in MCCs at stage 31. In control cells, we observed the stereotypical organization in apical and subapical networks that was previously described for mature MCCs (Werner et al., 2011). The apical actin network was organized like a grid surrounding each centriole, and the subapical network located just below was composed of short actin fibers that are known to connect neighboring BBs via their rootlets (Fig. 4A, A’). Both Lrrcc1 and Ccdc61 knockdown caused a global decrease of F-actin staining (Fig. 4D, E). When looking at individualized centrioles, a strong reduction of both apical and subapical F-actin was observed (Fig. 4B-C’). From these results, we conclude that Lrrcc1 and Ccdc61 are, either directly or indirectly, involved in the assembly of the apical filamentous actin networks.

**Figure 4.**
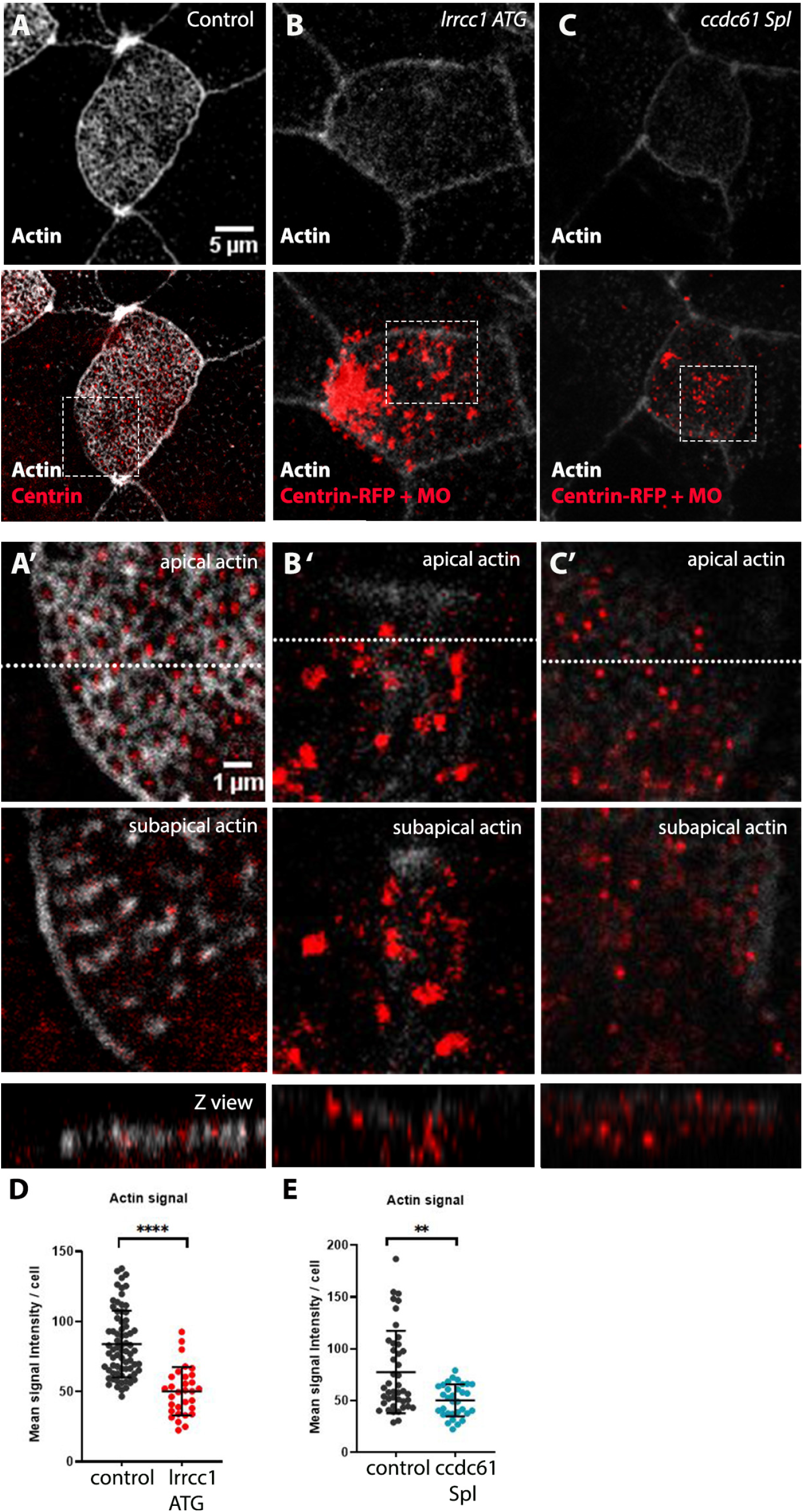
Depletion of Lrrcc1 and Ccdc61 impairs apical and subapical actin networks in MCCs. (A-C) MIPs of confocal acquisitions of St31 control and morphant MCCs stained for F-actin (white, Sir-actin), Centrin (red, Ab in control, Centrin-RFP co-injected with MOs to mark morphant MCCs). Dashed boxes indicate high magnification views in A’-C’. (A’-C’) Single slices of confocal acquisitions showing the apical or subapical (0.6 μm below) actin network in control and morphant cells. The dashed white line sets the position of the corresponding Z view (bottom row). (D-E) Graph displaying the quantification of mean F-actin signal intensity in control and morphant MCCs. Horizontal lines represent the mean and SD.

Next, we analyzed the apical MT network in mature MCCs. At stage 31, anti-α-Tubulin antibodies mainly revealed cilia, precluding the analysis of intracellular MT networks in mature MCCs (Fig. S7B). To circumvent this limitation, we adopted a deciliation strategy to deplete cilia-associated signals and visualize intracellular MTs (Fig. S7B). At stage 31, control cells were characterized by a highly organized apical MT network connecting BBs together (Fig. 5A, A’), similar to what has been reported by another method (Werner et al., 2011). In Lrrcc1 and Ccdc61 depleted-MCCs, intense α-Tubulin signals were found associated to clustered centrioles (Fig. 5B, B’, D, D’). When looking at individualized BBs, MTs were clearly visible but the size and spatial organization of filaments appeared very heterogeneous, as compared to control (Fig 5C, C’, E, E’). These results suggest that Lrrcc1 and Ccdc61 are not necessary for apical MT network polymerization *per se*, but could have a role, either directly or indirectly, on its organization.

**Figure 5.**
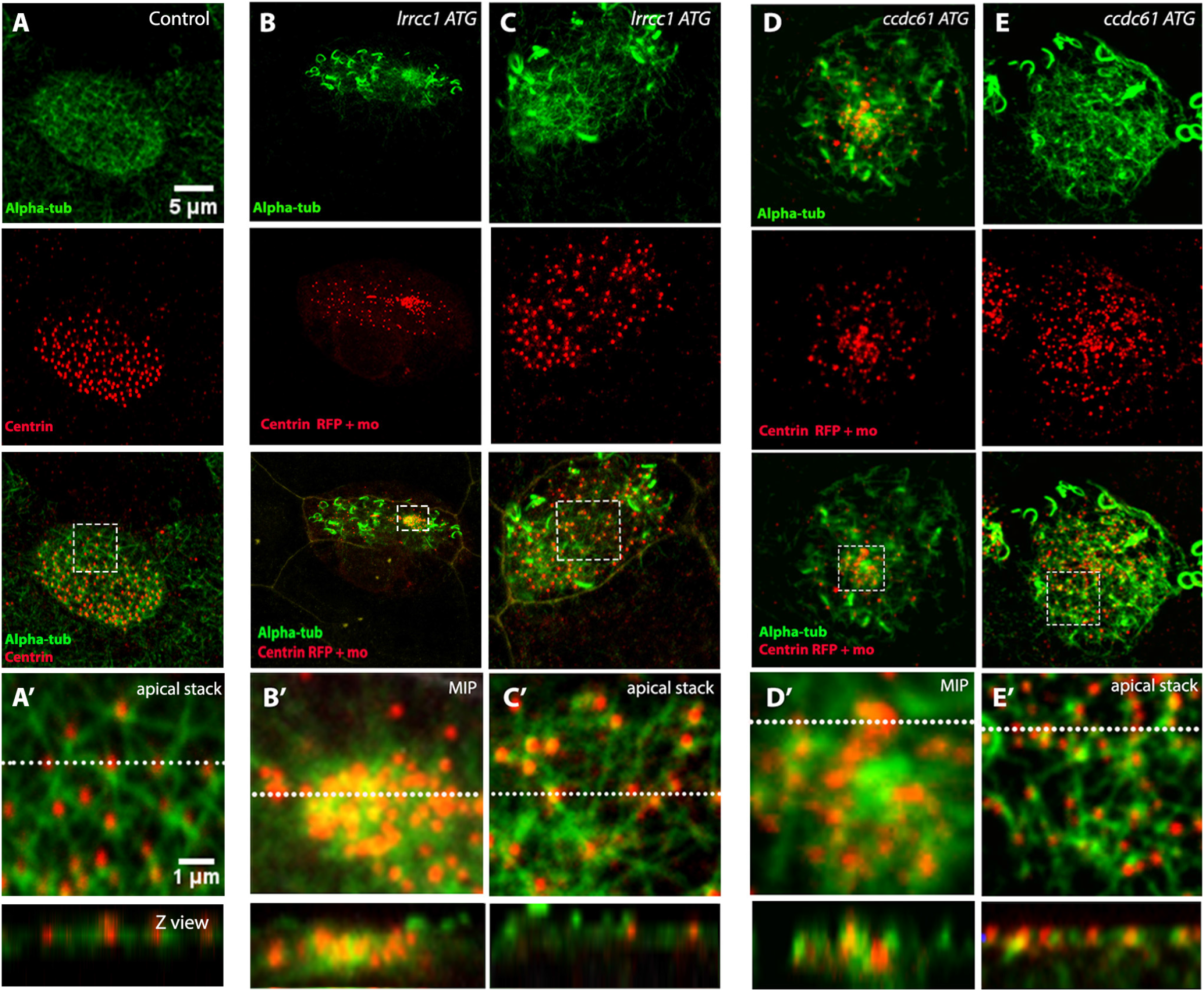
Depletion of Lrrcc1 and Ccdc61 causes apical MT network disorganization in MCCs. (A-E) MIPs of confocal acquisitions of St 31 control and morphant MCCs after deciliation stained with α-Tubulin (green, MTs) and Centrin Ab or expressing Centrin-RFP co-injected with MOs (red, BBs). White dashed boxes indicate the high magnification views in A’-E’. (A’, C’, E’) Apical confocal slices in control (A’), Lrrcc1 (C’) and Ccdc61 (E’) morphant cells. The regular MT network that links BBs in control cells appear irregular in morphant conditions. (B’, D’) MIPs of confocal acquisitions showing intense α-Tubulin signal around clustered BBs in Lrrcc1 and Ccdc61 morphant MCCs. Dashed white lines set the position of the corresponding Z views shown on the bottom row.

Finally, we analyzed the apical intermediate filament (IF) network using the anti-cytokeratin C-11 antibody. At stage 18, we did not detect IFs inside or at the MCC surface, suggesting that they are not involved in centriole apical migration (Fig. 6A). At stage 31, IFs were organized into a dense grid surrounding each BB (Fig. 6B, B’), similar to what has been described in tracheal MCCs (Tateishi et al., 2017). Strikingly, IF organization was drastically affected in Lrrcc1 and Ccdc61 morphant cells. Overall, the lattice appeared much less dense and the annular organization of IF around BBs was lost (Fig. 6C, C’, D, D’). This analysis suggests that Lrrcc1 and Ccdc61 are, directly or indirectly, involved in the establishment of the tight IF network in which BBs are embedded.

**Figure 6.**
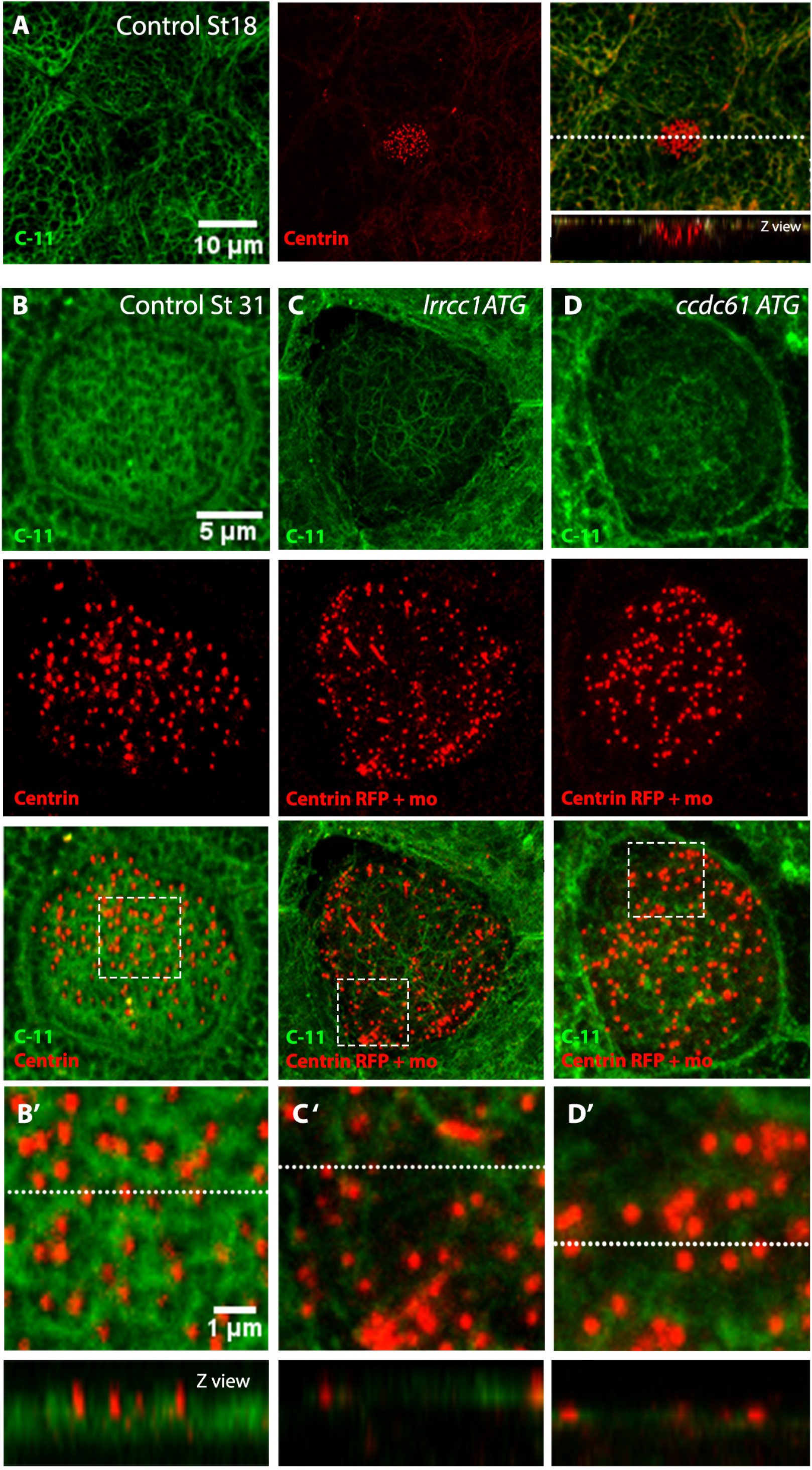
Depletion of Lrrcc1 and Ccdc61 impairs apical intermediate filament network in MCCs. (A-D) MIPs of confocal acquisitions of St 18 control MCC (A) and St 31 control (B) or morphant (C-D) MCC stained for IFs (C-11 Ab, green) and BBs (Centrin Ab or Centrin-RFP co-injected with MOs, red). Dashed boxes indicate high magnification views in B’-D’. (B’-D’) High magnification views of the zones boxed in B-D. The regular IF network that surrounds BBs in control cells appear much less dense in morphant conditions. White dashed lines set the position of the corresponding Z views shown on the bottom row.

Altogether these analyses reveal that Lrrcc1 and Ccdc61 depletion has profound impacts on MCC apical cytoskeleton organization.

### Lrrcc1 and Ccdc61 depletion reduces ciliary beating, impairs flow circulation and increases sensitivity to pathogen

The perturbed organization of BBs in morphant MCCs is expected to disturb the function of associated cilia, thereby affecting the production of fluid flow at the surface of the embryo. To address this issue, we first analyzed ciliary beating frequency by high-speed video-recording. As previously described, control MCCs performed synchronised and large amplitude effective and recovery strokes, characterized by the ability of cilia to extensively bend (Fig. 7A, and videos 1 and 2)(Werner et al., 2011). In Lrrcc1 and Ccdc61 morphant embryos, we observed three levels of beating defaults: (i) low-amplitude, uncoordinated and disoriented beating, causing occasional collisions between cilia; (ii) cilia performing only small vibrations; (iii) extreme cases with totally immobile cilia (Fig. 7A, and videos 1 and 2). Accordingly, the beating frequency, which was about 15Hz for the control condition, decreased to lower values in morphant MCCs. Around 20 and 30 % of MCCs were immotile in Lrrcc1 and Ccdc61 morphant conditions, respectively. In Lrrcc1 morphants, the remaining 80% of MCCs beat at much slower frequencies than control cells. In contrast, in Ccdc61 morphants, the remaining 70% of MCCs beat with similar frequencies as controls, suggesting that Ccdc61 depletion may cause an on/off switch (Fig. 7A). Next, we analyzed cilia-generated flow in the surrounding liquid, by live recording of visible dyed microspheres dispersed in the fluid, along the flanks of embryos at stage 31. In control condition, microspheres were moved by a robust flow and traveled the entire length of the embryo in approximately 12 seconds. In contrast, the flow was severely slowed down in Lrrcc1 and Ccdc61 morphant conditions, and the microspheres rarely reached the middle of the embryo after 12 seconds (Fig. 7B, C and video 3).

**Figure 7.**
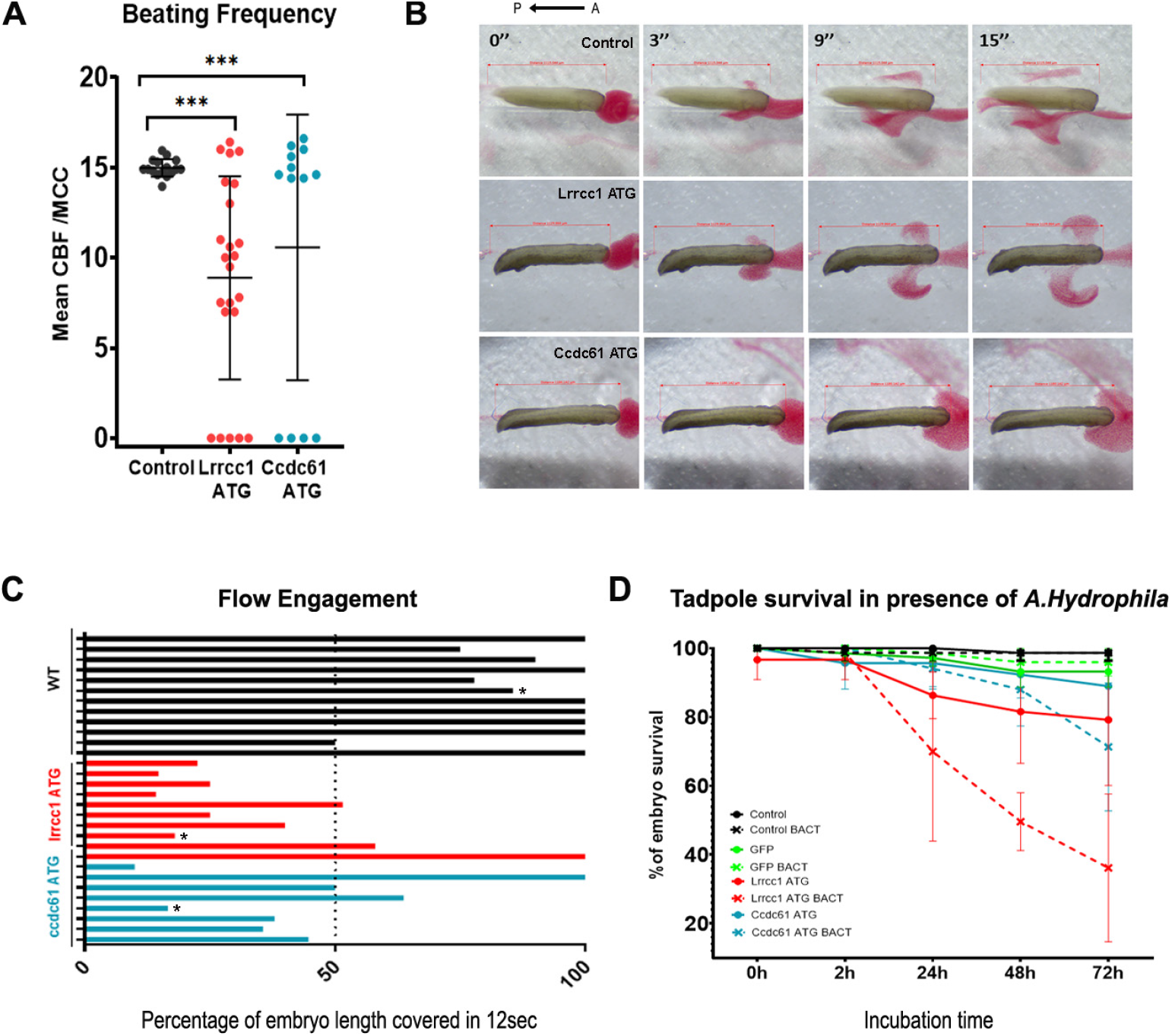
Lrrcc1 and Ccdc61 depletion impairs ciliary beating, flow production and embryo resistance against pathogens. (A) Quantification of cilia beating frequency (Hz) in control and morphant MCCs. Each dot represents the mean ciliary beat frequency computed over all visible cilia per individual MCC. Horizontal lines represent the mean and SD. (B) Still frames at 4 time-points taken from movie 3 showing the progression of the red dye along the flanks of control, Lrrcc1 and Ccdc61 morphant tadpoles. The black arrow represents the flow along the anterior-posterior axis (A-P). (C) Percentage of the embryo length reached by the dye front in 12 seconds. Each bar represents one recorded embryo. Cases marked with an asterisk are those shown in B. (D) Quantification of control and morphant tadpoles survival in presence or not of *A. hydrophila* bacteria.

To address the impact of impaired ciliary flow on the physiology of morphant individuals, we analyzed their susceptibility to pathogen infection. Embryos were incubated for 72h with the opportunistic bacteria *Aeromonas Hydrophila* (Dubaissi et al., 2018), and their survival rate was recorded. Non-injected and GFP-injected embryos were used as controls. For both control conditions, an almost complete survival rate was observed independently of the presence or not of bacteria (Fig. 7D). In absence of bacteria, Lrrcc1 and Ccdc61 morphant embryos displayed a slightly decreased survival rate as compared to control embryos. However, the presence of pathogenic bacteria strongly impacted the survival rate of morphants, starting from 24h of incubation. We confirmed by immunostaining that centrioles and cilia were disorganized in morphant embryos from the same experimental series (Fig. S8). This assay suggests that Lrrcc1 and Ccdc61 inactivation leads to a higher susceptibility of embryos to pathogen infection, likely due to reduced cilia-powered clearance.

## Discussion

In this study, we report a role for Lrrcc1 and Ccdc61 in *Xenopus* epidermal MCC differentiation and function, extending their evolutionary conserved importance in ciliated cells to vertebrates. At the cellular scale, both Lrrcc1 and Ccdc61 are necessary for the migration and/or docking of BBs at the cell surface and for apical cytoskeleton organization. At a larger scale, both factors are necessary for cilia-powered superficial flow to help survival of the organism, when exposed to environmental pathogens.

Lrrcc1 and Ccdc61 proteins both contain coiled-coil domains, which are one of the most common structural motifs mediating protein–protein interactions. Search in the IntAct protein interaction database reveals that Lrrcc1 and Ccdc61 physically interact with multiple known factors related to centrosomes, cilia and MT organization or polymerization, among which 30% are actually shared (Table 1)(Hein et al., 2015). Moreover, 20/26 (Lrrcc1), and 12/23 (Ccdc61) of those interactors were found to be expressed in MCCs of *Xenopus tropicalis*, as revealed by scRNA-seq (Table 1). Together with their reported interaction, this information is consistent with Lrrcc1 and Ccdc61 shared localization and phenotypes, and suggest that these two factors belong to a common molecular network important for BB function in MCCs.

**Table 1:** Lrrcc1 and Ccdc61 interactors. This table has been annotated from the Human Interactome Purification and Mass Spectrometry data of Hein et al., (2015). In green are shown common interactors between Lrrcc1 and Ccdc61. Enrichment values represent the average enrichment of the protein in multiples of standard deviations compared to control samples. Published evidence was screened to identify the existence or not of links with centrioles, cilia or MTs. Transcript expression in *Xenopus* tropicalis MCCs is displayed, as revealed by scRNA-sequencing (Briggs et al., 2018). Question marks correspond to transcripts that were totally absent from the available dataset.

| Gene name (bait) | Gene Name (prey) | Enrichment | Linked to centriole/cilia/MTs | Found in *Xenopus* MCCs |
| --- | --- | --- | --- | --- |
| LRRCC1 | LRRCC1 | 10,59121 | YES | YES |
| LRRCC1 | CCDC77 | 9,79952 | PROBABLE | YES |
| LRRCC1 | AZI1/CEP131 | 9,645587 | YES | ? |
| LRRCC1 | PCM1 | 8,385486 | YES | YES |
| LRRCC1 | CEP72 | 8,169336 | YES | YES |
| LRRCC1 | MIB1 | 6,578583 | YES | NO |
| LRRCC1 | FOPNL/FOR20/CEP20 | 6,448221 | YES | YES |
| LRRCC1 | BBS4 | 6,426498 | YES | YES |
| LRRCC1 | OFD1 | 6,392701 | YES | YES |
| LRRCC1 | WDR90 | 6,318789 | YES | YES |
| LRRCC1 | KIAA0753/OFIP | 6,241836 | YES | YES |
| LRRCC1 | CSPP1 | 6,139616 | YES | YES |
| LRRCC1 | KIAA1731/CEP295 | 5,889853 | YES | YES |
| LRRCC1 | PIBF1 | 5,812087 | YES | YES |
| LRRCC1 | CCDC18 | 5,503085 | PROBABLE | YES |
| LRRCC1 | WDR67 | 4,986441 | YES | ? |
| LRRCC1 | CCDC61 | 4,976773 | YES | YES |
| LRRCC1 | FGFR1OP/CEP43 | 4,853146 | YES | YES |
| LRRCC1 | SSX2IP | 4,807299 | YES | yes |
| LRRCC1 | CCDC14 | 4,215897 | YES | YES |
| LRRCC1 | KIAA1328 | 3,94653 | YES | NO |
| LRRCC1 | CEP350 | 3,833714 | YES | YES |
| LRRCC1 | PHLDB2 | 3,801527 | YES | NO |
| LRRCC1 | PRKAR2A | 3,785393 | YES | NO |
| LRRCC1 | TTLL5 | 3,725921 | YES | NO |
| LRRCC1 | N4BP3 | 3,684075 | NO | ? |
| LRRCC1 | CEP170 | 2,879607 | YES | YES |
| LRRCC1 | ESCO2 | 2,533443 | NO | ? |
| LRRCC1 | SPICE1 | 2,417896 | YES | YES |
| LRRCC1 | KIFC3 | 2,255614 | YES | NO |
| FGFR1OP | CCDC61 | 7,169243 | YES | YES |
| CEP120 | CCDC61 | 7,155219 | YES | YES |
| OFD1 | CCDC61 | 5,799062 | YES | YES |
| CEP72 | CCDC61 | 5,62468 | YES | YES |
| MIB1 | CCDC61 | 5,100073 | YES | NO |
| LRRCC1 | CCDC61 | 4,976773 | YES | YES |
| CEP135 | CCDC61 | 4,941259 | YES | YES |
| SDCCAG8 | CCDC61 | 4,656678 | YES | YES |
| CEP152 | CCDC61 | 4,148394 | YES | YES |
| KIF7 | CCDC61 | 3,749279 | YES | NO |
| TPX2 | CCDC61 | 3,734188 | YES | NO |
| NEDD1 | CCDC61 | 3,682512 | YES | YES |
| CEP55 | CCDC61 | 3,550385 | YES | NO |
| CCDC14 | CCDC61 | 3,546823 | YES | YES |
| SLAIN2 | CCDC61 | 3,535149 | YES | NO |
| PPP2CB | CCDC61 | 3,320352 | NO | NO |
| PCM1 | CCDC61 | 3,265138 | YES | YES |
| AZI1/CEP131 | CCDC61 | 2,932961 | YES | ? |
| CSNK1E | CCDC61 | 2,845102 | NO | NO |
| ANKRD26 | CCDC61 | 2,78639 | YES | YES |
| PRKAR2A | CCDC61 | 2,684373 | YES | NO |
| PPP2R5D | CCDC61 | 2,613387 | NO | ? |
| ZBTB33 | CCDC61 | 2,575864 | NO | NO |
| RAD54L | CCDC61 | 1,843647 | NO | NO |
| SRPR | CCDC61 | 1,821545 | NO | NO |

We show that Ccdc61 and Lrrcc1 associate to BBs early after their production and maintain this association in fully differentiated MCCs. The early association to neo-synthesized centrioles is compatible with a role in apical migration, which appears to be incomplete in Lrrcc1- and Ccdc61-deficient mature MCCs. Alternatively, apical migration may be unaffected, and defective docking may secondarily cause BBs to dive back into the cytoplasm. The distribution of Lrrcc1 and Ccdc61 at BBs is consistent with their observed requirement for correct MCC organization. This function is shared with Vfl1 (Lrrcc1) and Vfl3 (Ccdc61) orthologs, which are required to organize flagella/cilia in *Chlamydomonas*, *Paramecium* and planarians (Adams et al., 1985; Basquin et al., 2019; Bengueddach et al., 2017; Hoops et al., 1984; Wright et al., 1983). In contrast to studies in these species, however, we did not detect major rootlet structural defects upon Lrrcc1 and Ccdc61 depletion, suggesting the existence of redundant mechanisms to build up *Xenopus* BB appendages.

From their place of birth to their final position at the apical surface, the relocation of centrioles/BBs is intimately linked to cytoskeletal networks (Boisvieux-Ulrich et al., 1990; Herawati et al., 2016; Lemullois et al., 1988; Werner et al., 2011). The organization of these networks evolves during MCC maturation, allowing first to direct BBs towards the apical surface and then to orient and space them. By using chemical cytoskeleton inhibitors, it has been possible to alter migration, orientation or dispersion of BBs, thus attributing specific functions to actin filaments and MTs (Boisvieux-Ulrich et al., 1990; Werner et al., 2011). Our data revealed the primary importance of both Lrrcc1 and Ccdc61 for the establishment of actin filamentous networks. In particular, the absence of sub-apical actin fibers, which connect BBs through their rootlet (Antoniades et al., 2014; Werner et al., 2011), is consistent with the localization of Lrrcc1 and Ccdc61 at these sites. In contrast to actin filaments and MTs, the contribution of IFs to the organization of MCCs remains unknown. Our analysis did not reveal the presence of IFs at the stage of centriole apical migration, ruling out their implication at this step. In mature MCCs, a prominent IF network adopts an annular shape around BBs, similar to the apical F-actin network, but which appears to form more basally, extending below the BB level, where rootlets are found (Fig. 4A’ and 6B’)(Sandoz et al., 1988). This IF network collapsed in MCCs deprived of Lrrcc1 and Ccdc61, suggesting that IFs may stably interact with ciliary rootlets, through molecular complexes containing these two structural proteins. This first rudimentary analysis makes the *Xenopus* epidermis an attractive paradigm to address the specific role of IFs in MCC organization and function.

In contrast to the BF, which functions as an MT-organizing center (Clare et al., 2014; Kunimoto et al., 2012), the role of the rootlet appendage is less clear. Among proposed functions, it is generally believed to serve as an anchor, allowing BBs to maintain their position at the apical surface, and resist mechanical forces generated by ciliary beating (Antoniades et al., 2014; Bustamante-Marin et al., 2019; Yang et al., 2005; Yang and Li, 2005). In line with this, rootlets are scaffolds for molecular interactions, among which ciliary adhesion complexes were found to organize the short actin filaments that cross-link BBs together and help beating synchronization (Antoniades et al., 2014; Walentek et al., 2016; Werner et al., 2011). Here, we observed that Ccdc61, Pcnt and γ-Tubulin associate to rootlets in mature epidermal MCCs of *Xenopus.* Ccdc61 was recently shown to be a paralog of the scaffolding protein Sas6, known to establish the 9-fold rotational symmetry of MTs during centriole duplication (Ochi et al., 2020). Interestingly, an evolutionary conserved interaction between Sas6 and Pcnt has recently been reported (Ito et al., 2019). Furthermore, Pcnt is known for its ability to recruit γ-Tubulin to assemble a macro-molecular complex allowing MT nucleation at the centrosome, and at mitotic spindles (Woodruff et al., 2014). Based on this information and our own observations, it is tempting to propose that Ccdc61 may help Pcnt recruitment/stabilization at the rootlet, which in turn would favour γ-Tubulin recruitment and MT nucleation. Alternatively, the reported interaction of Ccdc61 with MTs (Ochi et al., 2020) may be important to link pillar MTs emanating from the BF (Clare et al., 2014) with nearby rootlets, which could further strengthen the mechanical coupling between adjacent BBs, to optimize their coordinated orientation and synchronized beating. Future super-resolution and EM studies should investigate the possible link between MTs and rootlets.

Unexpectedly, we found that cilia in Lrrcc1 and Ccdc61 morphant MCCs often beat very poorly. Proper ciliary beating entails correct axonemal structure and delivery of dynein motors to maintain ciliary motion (Huizar et al., 2018; Satir et al., 2014). Although we have not investigated these features, it is possible that the lack of Lrrcc1 and Ccdc61 compromises the capacity of rootlets to help trafficking towards the axoneme of essential structural or motor effectors (Gray et al., 2009; Mohan et al., 2013; Park et al., 2008; Yang and Li, 2005).

From the nanoscopic organization of organelles to the cellular function and physiology of the organism, all scales are coupled. The emergence of the locomotor function in planarians is a striking example of such coupling in multiciliated epithelia. In this model, polarization of BBs relies on their chiral construction, which mobilizes Lrrcc1 and Ccd61. This polarization allows the establishment of bilateral symmetry of the ventral multiciliated epidermis, which in turns governs the orientation of the worm movement (Basquin et al., 2019). In addition, retro-control loops exist between the different scales. For instance, in mouse ependymal MCCs, the establishment of a dense actin network that confers stability to BBs and cilia is dependent on active ciliary beating (Mahuzier et al., 2018). Thus, understanding such highly integrated systems implies analyzing multiple scales, as well as their coupling and feedback mechanisms. The present multi-scale study correlates the absence of structural proteins at BB appendages to MCC disorganization, defective flow production and impaired resistance to pathogens. Despite a wealth of studies on the *Xenopus* ciliated embryonic skin, it was unknown whether cilia-powered flow could limit infections by environmental pathogens. In that respect, the function of the *Xenopus* mucociliary epidermis is similar to that of the mucociliary epithelium that ensures the clearance of incoming pathogens in mammalian airways. In airways, cilia beating helps to propel a viscous layer of mucus trapping foreign particles, which lies on top of an aqueous periciliary layer. In contrast, in the *Xenopus* skin, the mucus layer is about 6 μm thick and sits right on top of the epithelium (Dubaissi et al., 2018), so that the 15 μm-long cilia actually beat in water to prevent attachment of micro-organisms. Thus, unlike in airways, the mucus on the frog embryonic skin may not be propelled as a coherent layer, which would explain the much lower density of MCCs necessary in this system. Although additional functions, such as oxygenation of the skin, are not ruled out, our proposed pathogen clearance function is consistent with the resorption of MCCs in pre-metamorphic tadpoles (Tasca et al., 2020), at a stage when innate immunity is in place (Robert and Ohta, 2009).

## Acknowledgements

Imaging in IBDM was performed on PiCSL-FBI core facility, supported by the French National Research Agency through the program “Investments for the Future” (France-BioImaging, ANR-10-INBS-04). The authors thank Florian Roguet for *Xenopus* care, Rémi Flores-Flores for ImageJ macro development, Brice Detailleur for 3D printing of embryo holders, Fabrice Richard, Nicolas Brouilly and Aicha Aouane from IBDM electron microscopy facility. This project was funded by grants from ANR (Oricen, 15-CE13-0003-02) and FRM (EQU201903007834). AN was supported by MESRI and FRM.

## Authors contribution

AN performed and analyzed most experiments. CB, OR, EB, and LK analyzed experiments. OR generated and characterized Pcnt antibodies. EB wrote scripts to analyze BB spacing. VT performed and analyzed Pcnt immunostaining. AV and EL designed, and EL performed and analyzed cilia beating experiments. AN and CB drafted the original manuscript. LK edited the manuscript. LK designed and supervised the study and obtained funding. All authors commented on the manuscript.

## Video legends

**Video 1: Examples of Lrrcc1 and Ccdc61 morphant MCCs with partially preserved ciliary beating**

Cilia beating was recorded at 250 fps, and the video is played at 50 fps to better appreciate cilia movement.

**Video 2: Examples of Lrrcc1 and Ccdc61 morphant MCCs without ciliary beating**

Cilia beating was recorded at 250 fps, and the video is played at 50 fps to better appreciate cilia movement.

**Video 3: Analysis of fluid flow in control, Lrrcc1 and Ccdc61 morphant embryos**

The video is played at real speed. Timing started when red microspheres were deposited onto the embryos.

## Supplementary figure legends

**Figure S1.**
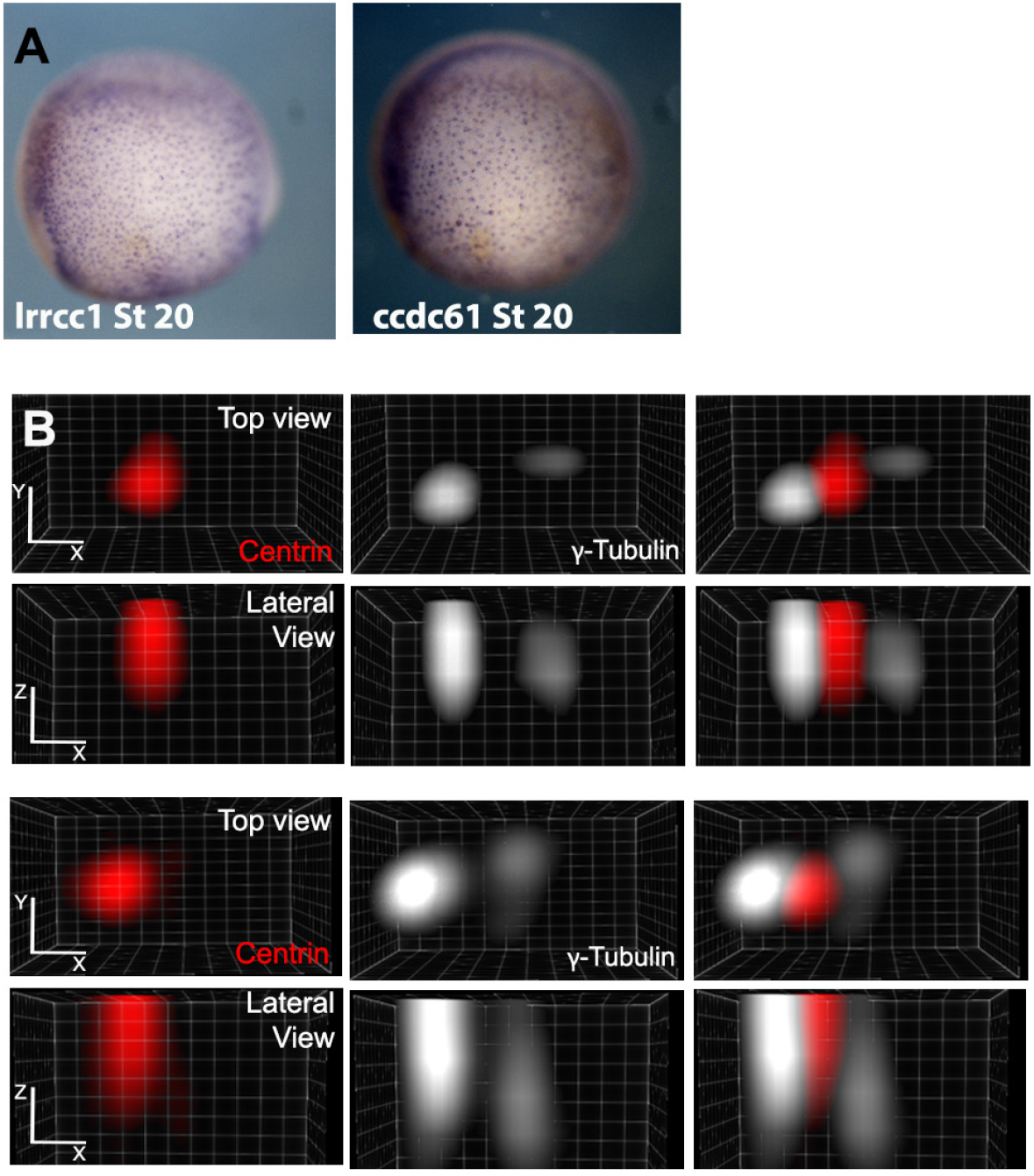
*lrrcc1* and *ccdc61* transcript expression in *Xenopus laevis* embryos. (A) *In-situ* hybridization for *lrrcc1* and *ccdc61* at St 20. (B) 3D Clear Volume top and lateral views of two individual BBs stained with γ-Tubulin (white) and Centrin (red) antibodies. γ-Tubulin protein localizes to BF and rootlets in *Xenopus* epidermal MCCs.

**Figure S2.**
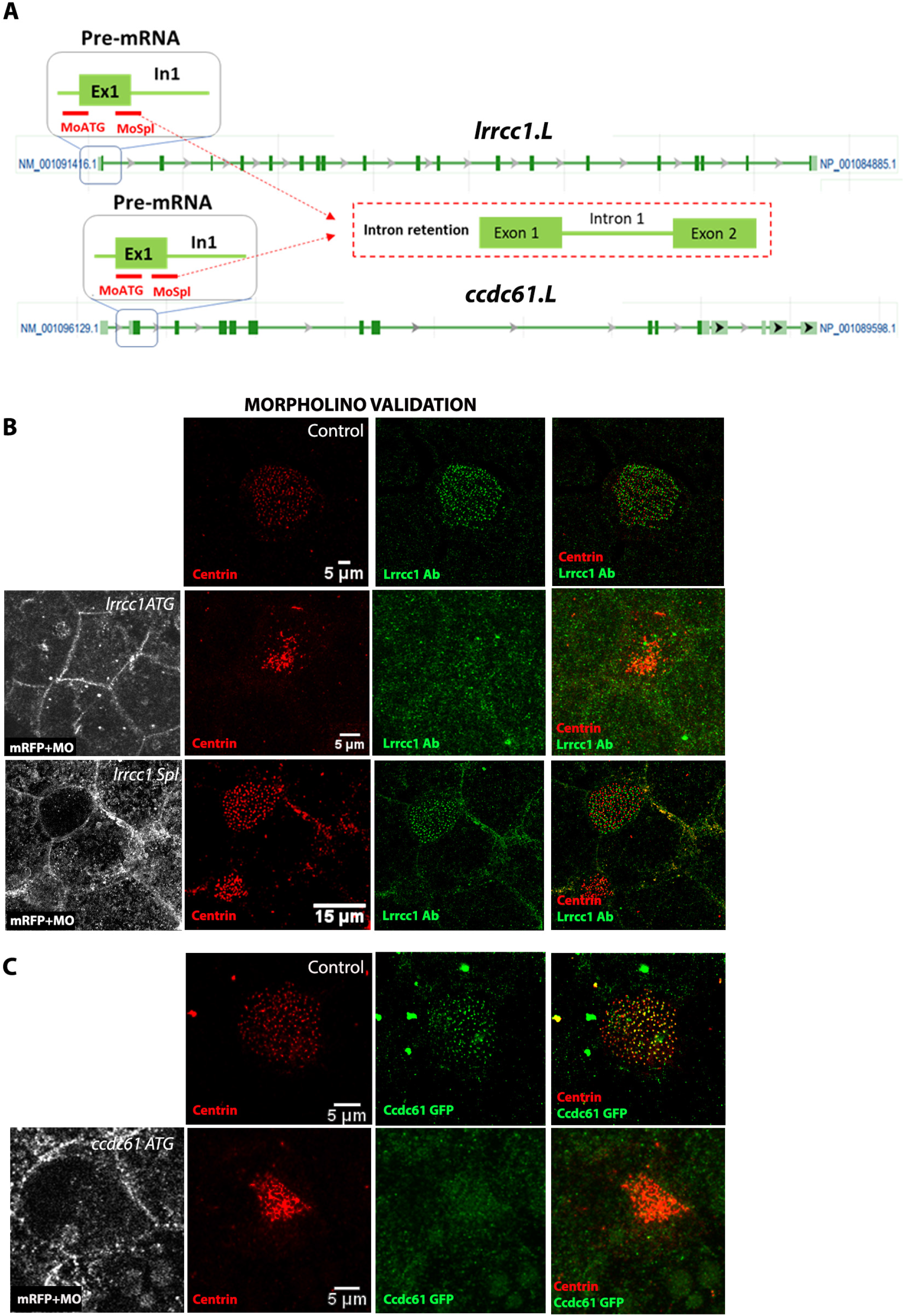
Lrrcc1 and Ccdc61 morpholino-mediated knock-down. (A) Schematic representation of *Xenopus lrrcc1* and *ccdc61* pre-mRNAs with introns and exons relative position and size. Red bars below exon1 show the position of *ccdc61* and *lrrcc1* MO ATG and Spl. (B) MIPs of confocal acquisitions of St31 control or morphant MCCs stained for Centrin (centrioles, red), Lrrcc1 (green) and mRFP (tracer co-injected with MO, white). MO efficiency is validated by disappearance of the Ab staining. (C) MIPs of confocal acquisitions of St 31 control or morphant MCCs expressing Ccdc61-GFP (green) and stained for Centrin (red) and mRFP (white). MO efficiency is validated by extinction of Ccdc61-GFP.

**Figure S3.**
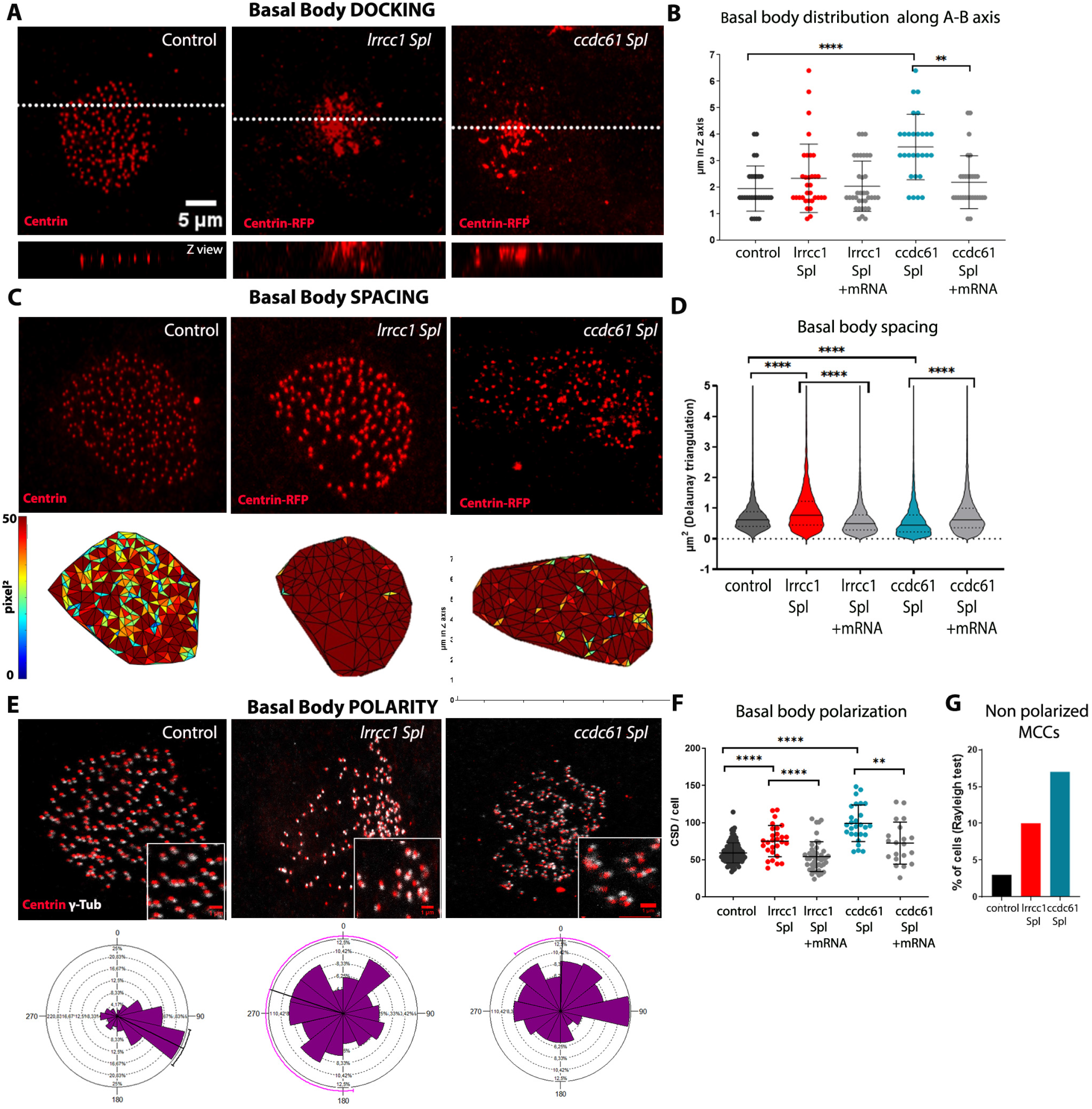
Lrrcc1 and Ccdc61 knockdown impairs BB docking, spacing and orientation. (A) MIPs of confocal acquisitions of MCCs from St 31 control, Lrrcc1 Spl or Ccdc61 Spl morphant embryos stained with Centrin Ab, or Centrin-RFP co-injected with MOs (red). Dashed lines set the position of the corresponding Z views presented below. (B) Graph displaying the number of Z slices containing BBs in control, Lrrcc1 and Ccdc61 morphant and rescued embryos. Each point represents a single cell. Horizontal lines represent the mean and SD. (C) MIPs of confocal acquisitions of apical BBs in MCCs from St 31 control, Lrrcc1 or Ccdc61 morphant embryos stained with Centrin Ab or Centrin-RFP co-injected with MOs (red) and Delaunay triangulation outputs (bottom row). Each colour is associated to a range of triangle areas in pixel^2^. (D) Violin plots displaying the distribution of triangle areas in μm^2^ of control, morphant and rescued MCCs. The horizontal line indicates the median, dashed lines indicate the quartiles. (E) MIPs of confocal acquisitions of control and morphant MCCs stained with Centrin (red, centrioles) and γ-Tubulin (white, BF). Below each cell, its respective rose histogram representing the distribution of BB orientations. The black line running from the centre of the diagram to the outer edge is the mean angle and the arcs extending to either side represent the confidence limits of the mean (when the mean angle is only theoretical but not significant, the line turns pink). (F) Graph displaying the percentage of non-polarized MCCs (no significant mean angle of BBs within a cell can be calculated), as revealed by the Rayleigh statistical test. (G) Circular Standard Deviations of control, morphant and rescued MCCs. Each point represents a single cell. Horizontal lines represent the mean and SD. All confocal images are at the scale shown in (A).

**Figure S4.**
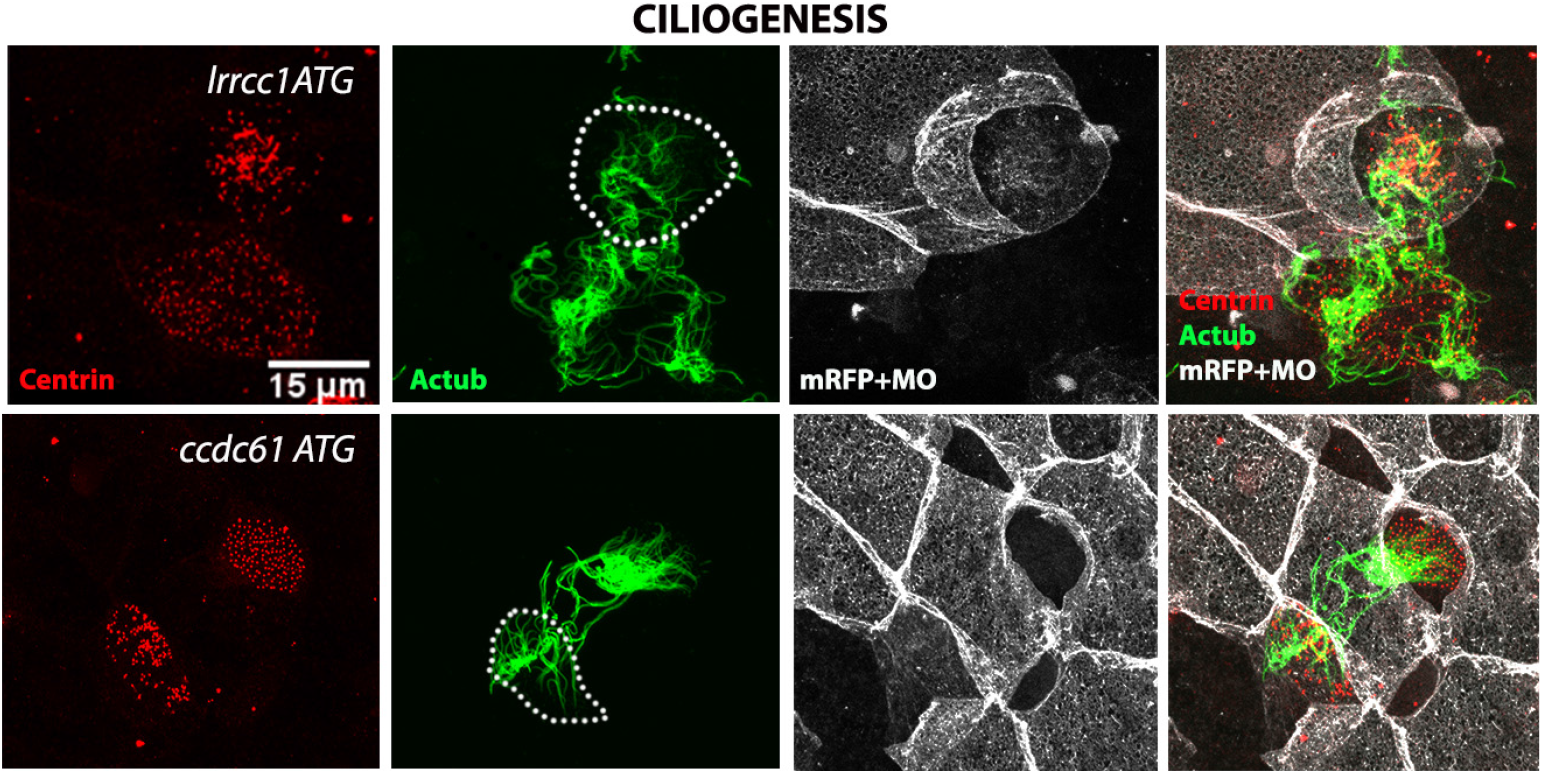
Ciliogenesis is not affected by Lrrcc1 and Ccdc61 depletion. MIPs of confocal acquisitions of St 31 control and morphant MCCs stained for centrioles (Centrin Ab, red), cilia (Acetylated-α-tubulin Ab, green) and morpholino tracer (mRFP co-injected with MOs, white). White dashed lines indicate morphant MCCs.

**Figure S5.**
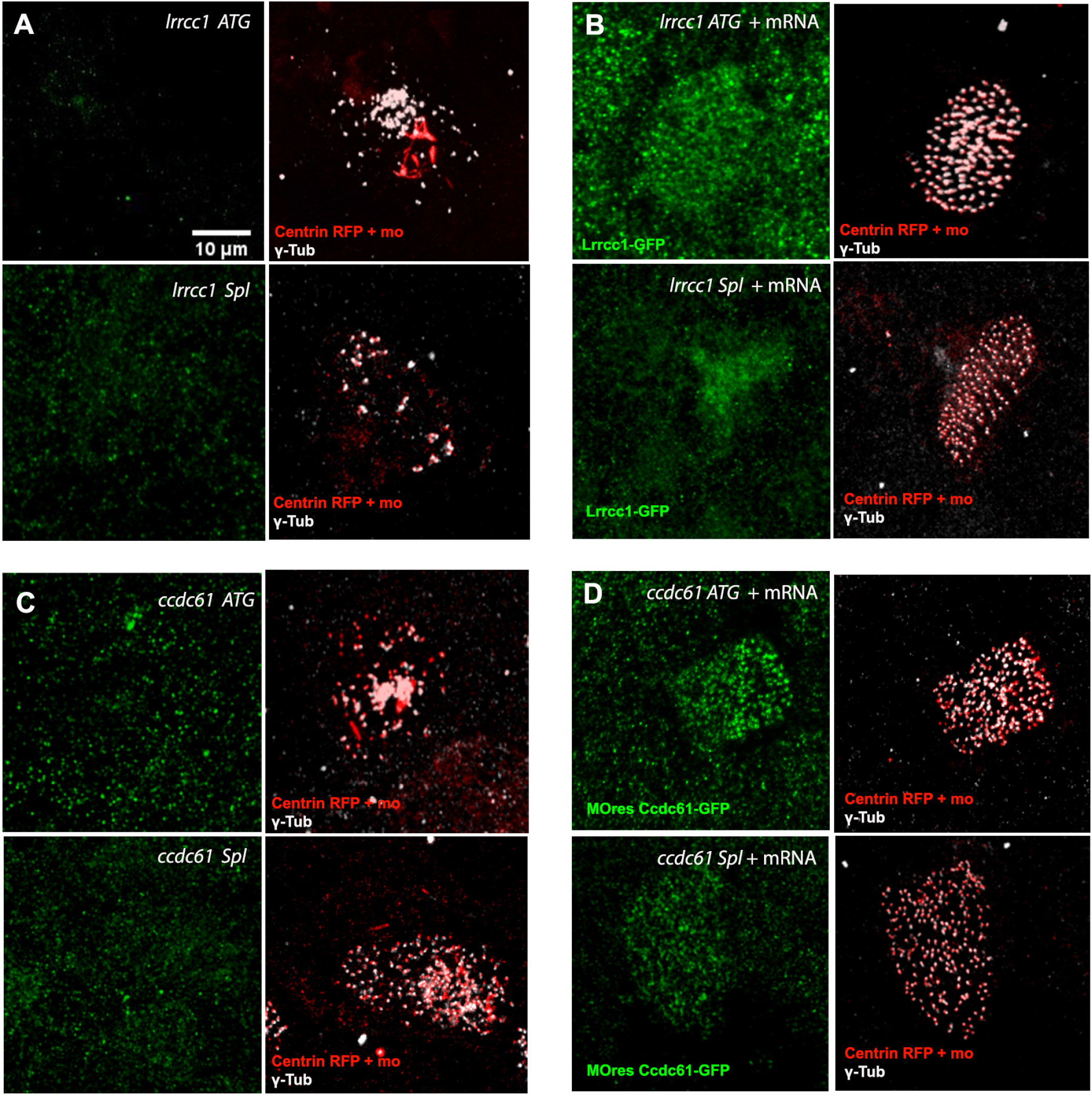
Correct BB organization is rescued upon Lrrcc1-GFP and Ccdc61-GFP mRNA co-injection in morphant MCCs. (A-D) Embryos were sequentially injected first with MO + Centrin-RFP and then with Lrrcc1-GFP or MOresCcdc61-GFP mRNA (green). (A, C) MIPs of confocal acquisitions of St 31 MCCs expressing Centrin-RFP (BBs, tracer for MO injection, red), immunostained for BF (γ-tubulin, white). The absence of specific signal in the green channel identifies those cells as morphant MCCs, consistent with severe BB disorganization. (B, D) MIPs of confocal acquisitions of MCCs expressing Centrin-RFP co-injected with MOs (red), Lrrcc1-GFP or MOresCcdc61-GFP (green) and immunostained for BF (white). In GFP-positive morphant MCCs, BB organization is largely rescued and comparable to control (see Fig. 2 and S3).

**Figure S6.**
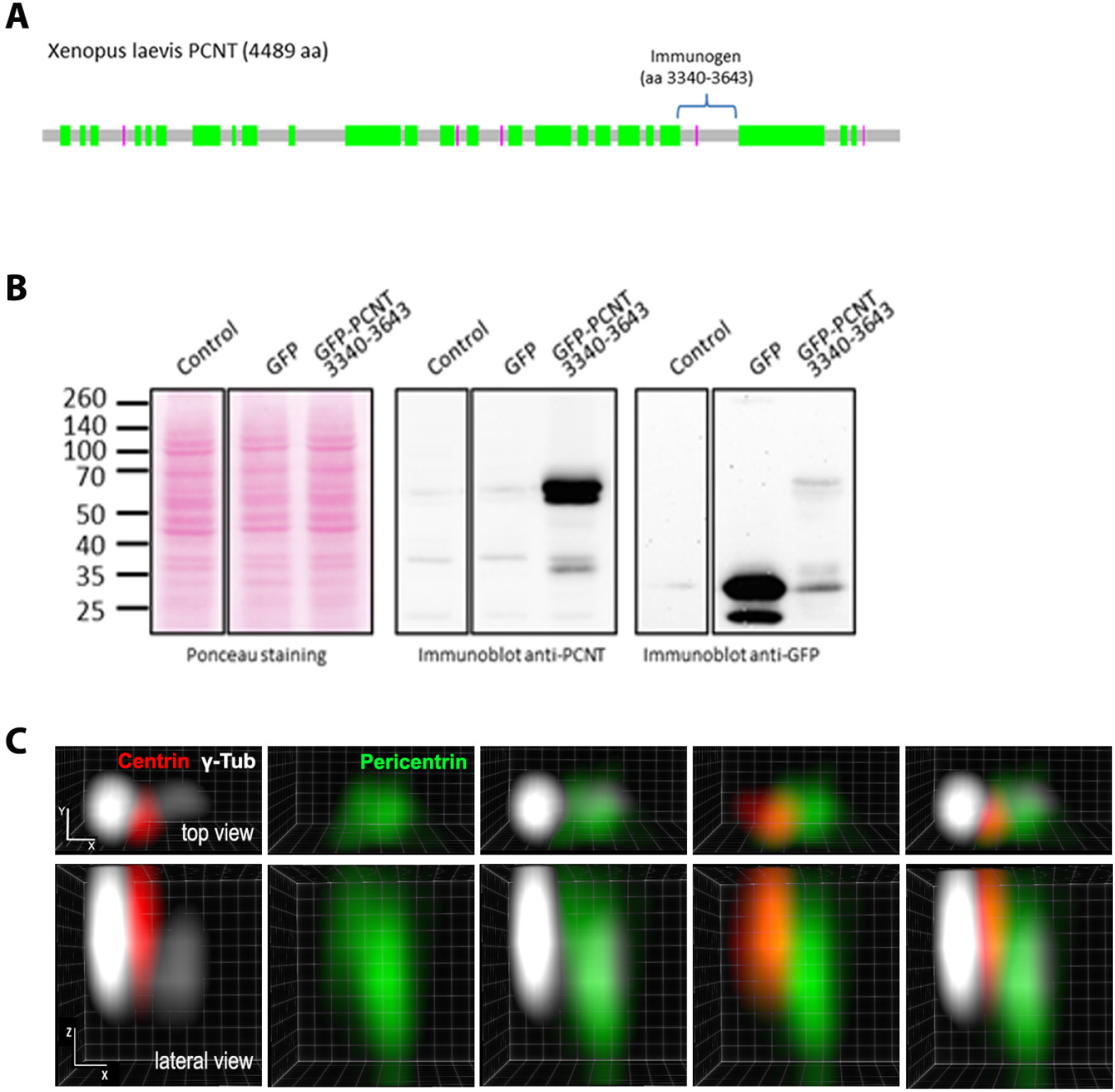
Pericentrin antibody validation. (A) Schematic representation of *Xenopus laevis* Pcnt protein. A portion of the protein between aa 3340-3643 was used as immunogen to generate rabbit polyclonal Ab. (B) COS1 cells were transfected or not (control) with vectors coding for the indicated proteins and immunoblotted with custom-made Pcnt rabbit Ab, or Ab against GFP. The expected size of GFP-Pcnt is 62 kDa. (C) Clear Volume 3D top and lateral views of a single immunostained BB, showing the localization of Pcnt (green), relative to Centrin (red), and γ-Tubulin (white).

**Figure S7.**
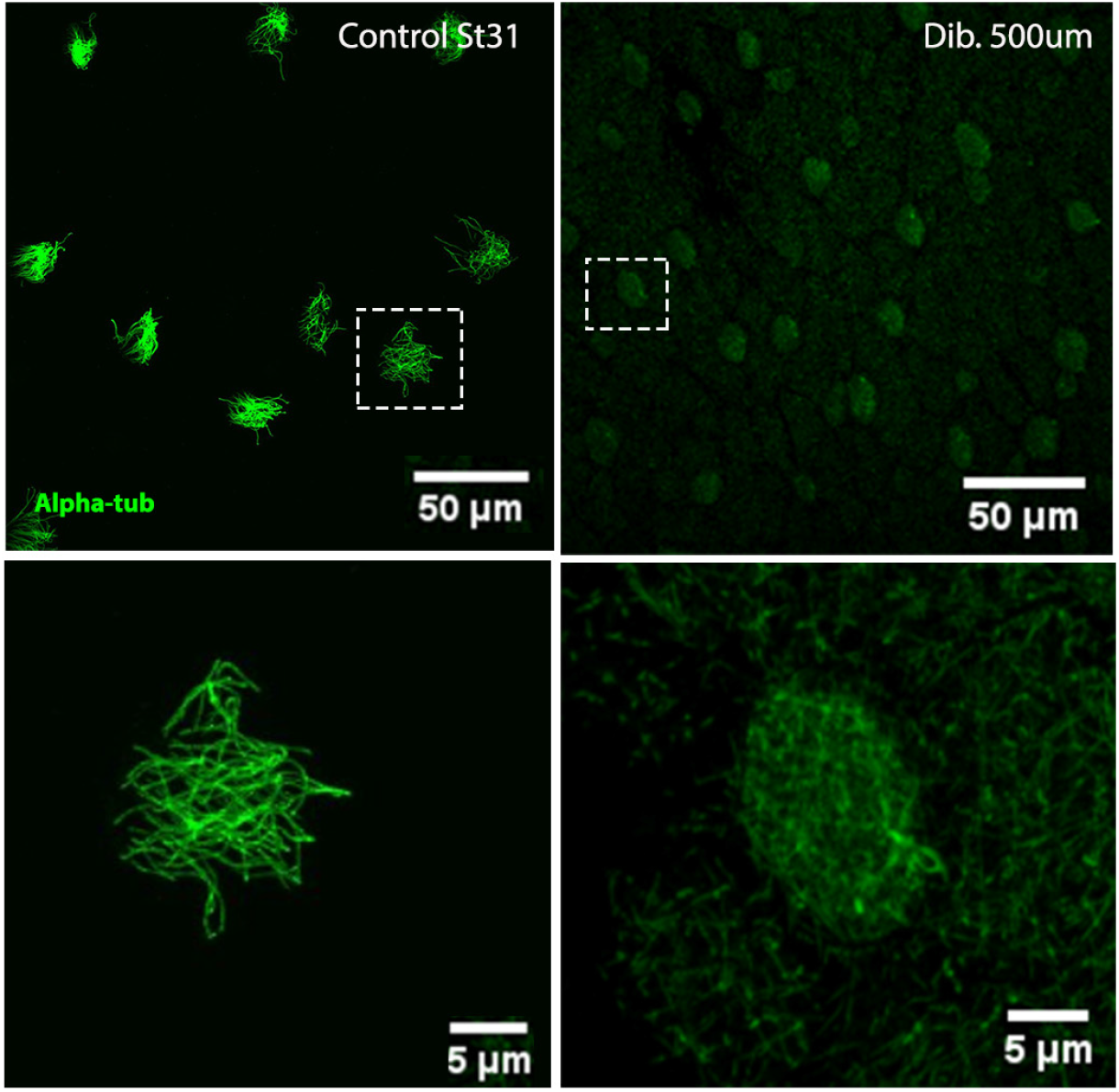
Deciliation of *Xenopus* epidermal MCCs. MIPs of confocal acquisitions of the ciliated epidermis before and after dibucaine treatment. Cilia and MTs are stained with α-Tubulin Ab (green). The absence of cilia upon dibucaine treatment allows the observation of cytoplasmic MTs.

**Figure S8.**
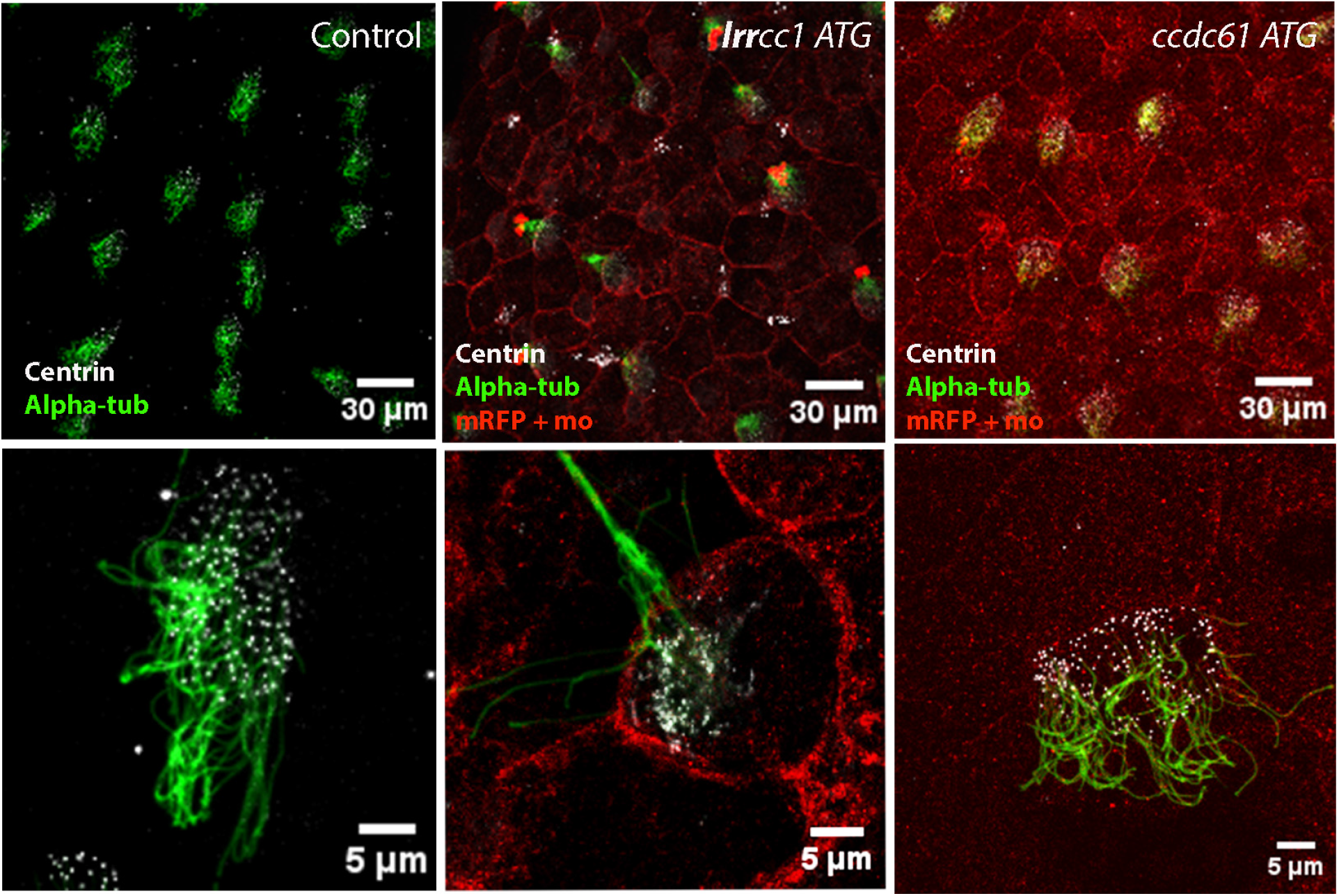
Phenotype validation of embryos incubated with *A. hydrophila*. MIPs of confocal acquisitions of epidermis of St 31 embryos, from the same experimental pool as those used for our survival assay (Fig. 7), stained for cilia (α-Tubulin Ab, green), centrioles (Centrin Ab, white) and MO tracer (mRFP Ab, red). The visible BB disorganization confirms MO efficiency.

## Materials and methods

### Ethics statement

All experiments were performed following the Directive 2010/63/EU of the European parliament and of the council of 22 September 2010 on the protection of animals used for scientific purposes and approved by the “Direction départementale de la Protection des Populations, Pôle Alimentation, Santé Animale, Environnement, des Bouches du Rhône” (agreement number F1305521).

### RNA probes and whole mount *in situ* hybridization

cDNA fragments from *Xenopus laevis lrrcc1*.L (entrez gene 431936), and *ccdc61*.L (entrez gene 734655) were amplified from commercial cDNA (Horizon discovery) by PCR using the following primers:

*lrrcc1* forward: 5′-GCGAACGGACACAGACAGTA-3′;

*lrrcc1* reverse: 5′-GAATTCCATGGTAGTCAGCTCCTGC-3′;

*ccdc61* forward: 5′-GCGGCCGCAAGTGGAGGATGCTGTGACC-3′;

*ccdc61* reverse: 5′-GAATTCACGGATGAACTGCGTCTCTG-3′.

PCR products were cloned in pBlueScript KS+ vector and digoxigenin-labelled probes were generated from linearized plasmids using RNA-labeling mix (Roche). Whole-mount chromogenic *in situ* hybridization was performed as described previously (Marchal 2009) using 40ng of Digoxygenin-labelled probe. Pictures were taken with the stereomicroscope Leica MZ125 coupled to NIKON digital Sight DS-Fi1 camera.

### Plasmids, RNAs, and Morpholinos

To generate plasmids for RNA synthesis and micro-injection, the ORFs of *lrrcc1* and *ccdc61* were amplified by PCR using the following primers:

*lrrcc1-*forward: 5′ TCTTTTTGCAGGATCACAATGGCAGGCACGGACCCACGAA 3′;

*lrrcc1-*reverse: 5′ CTTTACTCATTCTAGAAAATTCTTTTTGGATGTCACTTAG 3′;

*ccdc61-*forward: 5′ TCTTTTTGCAGGATCACAATGGAGGATACAGAGTTTGCT 3′;

*ccdc61-*reverse: 5′ CTTTACTCATTCTAGACTGCATCAGTAAGTACCCGCTGGCT 3′.

PCR products were subcloned in frame with GFP sequence in 3′ into pCS2+-GFP vector using In-Fusion^®^ HD Cloning Kit (Takara Bio USA, Inc.). For rescue experiments, silent mutations were introduced by PCR in the original Ccdc61 sequence (3′ GAG-GAT-ACA-GAG-TTT-GCT-GAA-G 5′) to generate a Ccdc61-GFP construct (named MOresCcdc61-GFP) with a sequence resistant to MO ATG (3′ GA**A**-GA**C**-AC**G**-GA**A**-TT**C**-GCT-GAA 5′). pCS2+-mRFP was used to generate an injection reporter.

The cDNA fragment coding for amino-acids 3340 to 3643 of the full-length *Xenopus laevis* Pcnt was amplified by PCR from a partial cDNA clone (IMAGE 5156155, Source Bioscience). PCR products were cloned by Gateway^®^ recombination into pGEX6P3 (GE Healthcare Life Sciences) and pEGFP-C1 (Clontech) to produce GST fusion proteins and express GFP-tagged proteins, respectively. The Centrin-RFP plasmid was a kind gift from JB Wallingford.

Capped mRNAs were synthetized from linearized vectors using the SP6 mMESSAGE mMACHINE^®^ Kit (Ambion Life Technologies) and purified with the MEGAclear™ Kit (Ambion Life Technologies).

Two independent morpholino antisense oligonucleotides were designed against *lrrcc1* and *ccdc61* (GeneTools, LLC). *lrrcc1*-ATG-MO: 5′ GTGCCTGCCATTCTCCCGCAACAAA 3′; *lrrcc1*-Spl-MO: 5′ ACTGAAGCCATGCTGCTTACCTGGA 3′; *ccdc61*-ATG-MO: 5′CTTCAGCAAACTCTGTATCCTCCAT 3′; *ccdc61*-Spl-MO: 5′TGTCTCCCACTTCTACTCACATTGA 3′.

### *Xenopus* embryo injections

Eggs obtained from NASCO females were fertilized in vitro, dejellied and cultured as described previously (Marchal et al., 2009). 20-30 ng of MO was injected (alone or with 200-500pg of mRFP tracer) in one or two animal-ventral blastomere (presumptive epidermis) at different stages depending on the experiment (Table 2). Working amounts of MOs were first calibrated to optimize embryo survival. Validation of MOs efficacy in depleting the target protein was confirmed based on Lrrcc1 antibody and Ccdc61-GFP signal disappearance. For rescue experiments, a sequential injection strategy was adopted to obtain mosaic embryos containing differentially marked morphant and rescued MCCs. At 4-cell stage*, lrrcc1* and *ccdc61*-ATG-MOs + Centrin-RFP or *lrrcc1* and *ccdc61*-Spl-MOs + Centrin-RFP was injected At 16-cell stage, the same embryos were injected with Lrrcc1-GFP or MOresCcdc61-GFP mRNAs. After injection, embryos were incubated at different temperatures between 13°, 18° and 23°C in MBS 0.1x until they reached the desired developmental stage. Whole-embryos were fixed in different conditions (Table 3) and stored in 100% methanol at −20°C in preparation for immunofluorescent staining.

**Table 2:**
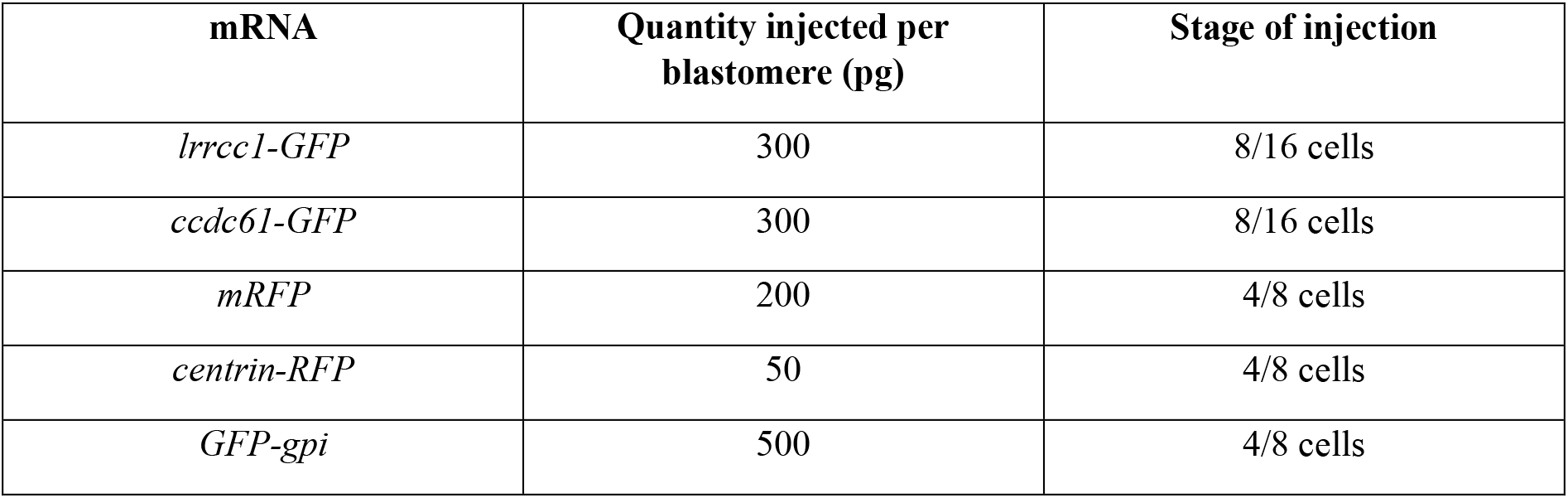
mRNA injection conditions.

**Table 3:** Primary antibodies and markers.

| Fixation | Target/Species/Isotype | Reference | Dilution |
| --- | --- | --- | --- |
| MetOH 100% 4°C<br>OverWeekend or<br>PFA 4% 0,1%<br>Triton 30min RT | γ-Tubulin mouse IgG1 | Abcam Ab11316 | 1/800 |
|  | Acetylated-tubulin mouse IgG2b | Sigma-Aldrich T7451 | 1/1000 |
|  | GFP chicken | AvesLab GFP-1020 | 1/500 |
|  | RFP rat IgG2a | Chromotek 5F8 | 1/500 |
|  | Centrin mouse IgG2a | Sigma-Aldrich 04-1624 | 1/1000 |
|  | Cytokeratin (Pan-reactive) mouse monoclonal IgG1 | Sigma-Aldrich SAB4700666 | 1/500 |
|  | Z01 mouse IgG1 | Thermo Scientifique 33-9100 | 1/200 |
| | α-Tubulin mouse IgG1 | Sigma-Aldrich T9026 | 1/500 |
| Memfa 1h RT | Sir-Actin | Spirochrome SC001 | 1/1000 |
| Memfa 2h RT or OverNight 4°C | Anti-DIG-PA conjugated IgG | Roche 11266026 | 1/500 |
| PFA 4% Triton 0,1x 30min RT | Pericentrin rabbit | Home made, this study | 1/5000 |
| MetOH 100% 4°C OverWeekend | Lrrcc1 rabbit polyclonal | Sigma-Aldrich HPA012893 | 1/1000 |

### Embryo deciliation

Stage 31 embryos were incubated for 2h in a 35-mm Petri dish containing 2 ml of dibucaine hydrochloride (200μM in MBS 0.1X; Fluka D0638-16). After a rapid wash in MBS 0.1x embryos were immediately fixed by incubation in MetOH 100% at −20°C for two days before being further processed for immunofluorescent staining.

### Immunostaining

After a sequential re-hydration in solutions with decreasing methanol concentrations, embryos were incubated in blocking solution (3% Bovine Serum Albumin (BSA) in PBS 1x) for 1h at room temperature (RT) and subsequently in primary antibodies diluted in 3% BSA overnight at 4°C (Table 2). After washing in PBS 1x embryos were incubated with fluorescently labelled secondary antibodies diluted in 3% BSA for 1h at RT (Table 4). After washing, embryos were mounted in Mowiol (Sigma-Aldrich) between slide and coverslip.

**Table 4:**
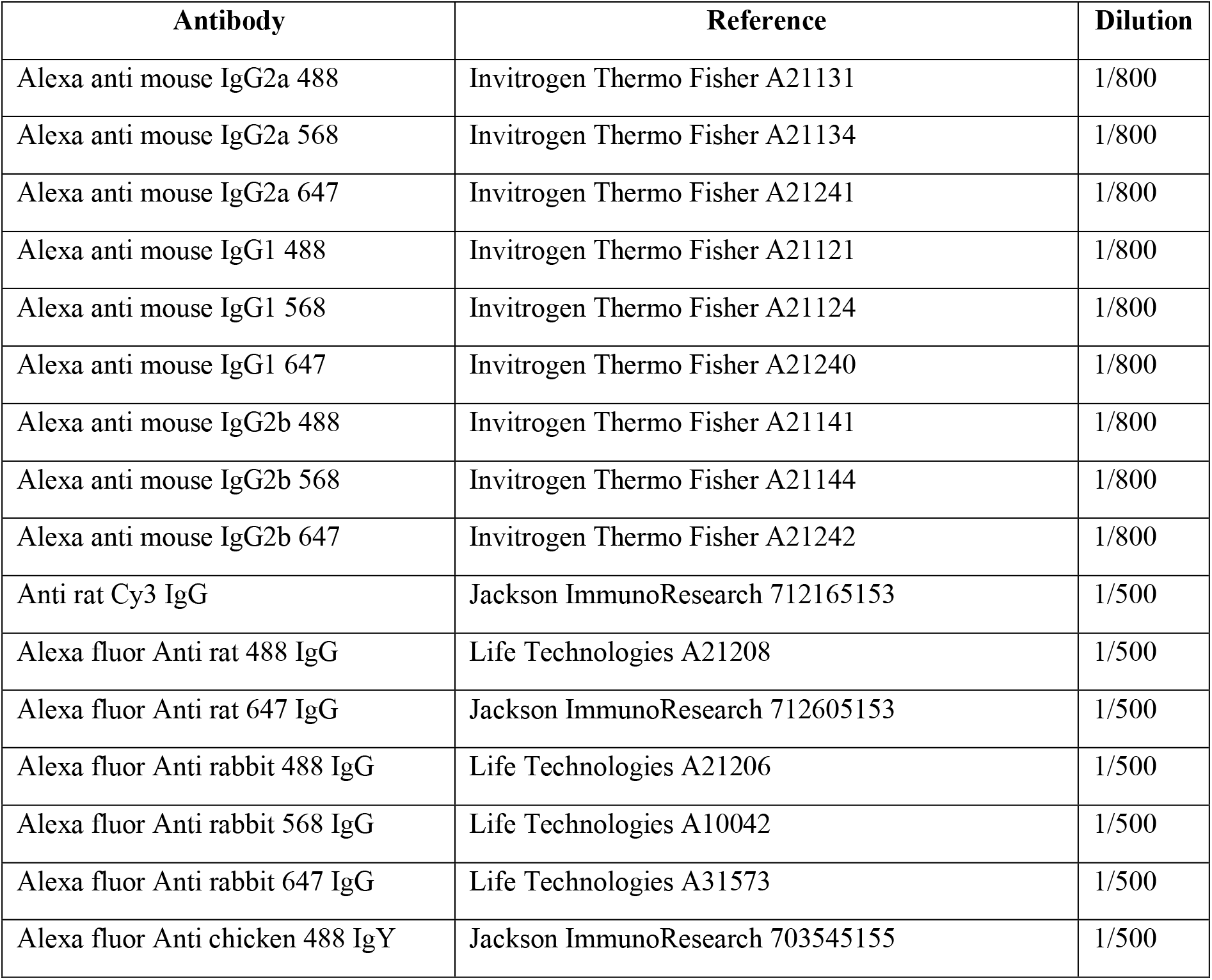
Secondary antibodies.

### Confocal Microscopy

Confocal pictures were acquired using ZEISS LSM 780 right standing AxioImager Z2 and ZEISS LSM 880 reverse standing AxioObserver 7 equipped with 20x and 63x oil objectives. Two- or three-colors confocal z-series images (z-slices interval between 0.3-0.8nm) were acquired using sequential laser excitation. When necessary, images were converted into single plane maximum intensity projection (MIP) and edited using Image J 2.0.0 software.

### Image analysis

F-actin and Pcnt signals were quantified by measuring the mean pixel intensity per MCC with ImageJ.

BB docking was quantified manually by counting the number of z-slices containing Centrin- or Centrin-RFP-positive centrioles along the apical-basal axis for each cell. These values were transformed in μm based on the interval size between slices set during confocal acquisition. BB spacing analysis was performed from Centrin or Centrin-RFP immunostaining images using custom made Matlab scripts. After manual segmentation of individual cells, the script (i) automatically detects, and segment individual BBs within each cell. Only apically docked and isolated BBs are considered, omitting those located below the apical cell membrane or in dense clusters; (ii) determine the centroid of each BB; (iii) use XY coordinates of all the centroids to build triangles between the nearest neighbors with the Delaunay Triangulation Matlab script. Once the triangulation is obtained, areas of the triangles are measured in pixel^2^ and transformed in μm^2^ based on the pixel size of each acquisition.

BB orientation was analysed using a home-made ImageJ script (designed by R. Flores-Flores). The script (i) automatically detects individual MCCs in the field of view; (ii) automatically detects individual BBs in each MCC; (iii) traces a vector from Centrin to γ-Tubulin spots (a threshold is applied for γ-Tubulin channel in order to detect only the BF associated spot, and not the rootlet spots) for each BB and calculate its angle with the vertical axis. The output of the script is a list of angles that were plotted using the Oriana software (Version 4.02, Kovach Computing Services) to obtain a graphical representation of their distribution in 25,7° bins, with 95% of confidence intervals. Finally, circular statistical analysis were performed (CSD, Rayleigh’s Uniformity Test).

### Transmission electron microscopy

Embryos were processed for electron microscopy as previously described (Revinski 2018) and cut transversely at midbody level (80 nm/slice) with a Leica Ultracut UC7 (Leica, Germany). Images were acquired using a Tecnai G2 (Thermofisher, USA) microscope equipped with a Veleta camera (Olympus, Japan).

### Cell culture and Western Blot

Cos-1 cells were grown in DMEM supplemented with 10% heat inactivated FCS and transfected with Fugene HD (Roche Applied Science) according to manufacturer’s protocol. Transfected or control cells were washed in PBS and lysed in 50 mM Tris HCl pH 7.5, 150 mM NaCl, 1 mM EDTA, containing 1% NP-40 and 0.25% sodium deoxycholate (modified RIPA) plus a Complete Protease Inhibitor Cocktail (Roche Applied Science) on ice. Cell extracts separated on polyacrylamide gels were transferred onto Optitran membrane (Whatman), followed by incubation with rabbit anti-GFP (1:5000, Abcam, ab290), or homemade rabbit anti-Pcnt antibody (1 μg/ml) produced by immunization with recombinant portion of *Xenopus laevis* Pcnt (XP_018091513.1, residues 3340-3643), and horseradish peroxidase-conjugated secondary antibody (Jackson Immunoresearch Laboratories, 711-035-152). Signal obtained from enhanced chemiluminescence (Western Lightning ECL Pro, Perkin Elmer) was detected with MyECL Imager (ThermoFisher Scientific).

### Flow and cilia beating measurements

Stage 31 living embryos were placed in anaesthetic solution media (MS222 0.02% in MBS 0.1x) in a home-made 3D-printed Petri Dish specially designed to maintain them with the dorsal side on the top. The setup was placed under compact Stereo Microscope ZEISS Stemi 305 coupled to ZEISS Axiocam 105 Color microscope camera. Timelapse recording was done after release of 1μl of visible dyed microspheres (Bangs Laboratories: DSCR006, Mean diameter 5.19 μm, diluted 1/2 in MBS 0,1x) at the anterior front of the embryo.

Cilia beating frequency was analysed on the same pool of living embryos. Embryos were placed between a glass slide and a coverslip with a drop of anaesthesic medium (MBS 0.1x – MS222 0.02x) surrounded by grease (high vacuum grease DOW CORNING, Sigma-Aldrich Z273554-1EA) to stick the coverslip. Movies with a duration of 3 seconds (250 fps) were recorded at the ventral border of the embryos with a Nikon eclipse Ti-E microscope and a 63x long working distance air objective. Computation of cilia beat frequency (CBF) was done using an in-house routine developed in python (Khelloufi et al. 2018). The resulting frequencies (Hz) shown in the quantification represent the mean CBF computed over all visible cilia per individual MCC. Movies are played at 50fps to observe better the beating strokes and defaults. After flow and CBF recording, MO efficiency was assessed in those embryos by immunostaining for Centrin, Acetylated-α-tubulin and mRFP (injection tracer).

### Embryo survival to *Aeromonas hydrophila*

*Aeromonas hydrophila* (Chester Stanier, ATCC®7966 kindly provided by E. Dubaissi) colonies were grown for 48h at 37°C on LB Agar + 30μg/mL Kanamycin. Next, individual colonies were cultured in 10mL LB + 30μg/mL Kanamycin overnight at 37°C. The OD_600_ (OD600 DiluPhotometer version1.4, Implen) was measured and the culture was either diluted in LB or grown further to reach an OD_600_ of 0.2. Cultures were centrifuged 10 min at 3500 xg and the bacteria pellet was resuspended in the same volume of MBS 0.1x. Controls (Non-injected or mRFP-injected) and morphant (Lrrcc1-MO+mRFP or Ccdc61-MO+mRFP) embryos were incubated from St 31 during 72h at 13°C in 3mL of bacteria containing medium. Survival was assessed by recording active response to touch at 2h, 4h, 24h, 48h and 72h. At the end of the experiment, MO efficiency was assessed by immunostaining for Centrin, Acetylated-α-tubulin and mRFP tracer.

### Statistics

For all experiments, statistical analysis of significance was done using GraphPad Prism (version 8.2.0 for Windows, GraphPad Software, San Diego, California USA). First, the normality of data (Gaussian distribution) was tested using D’agostino & Pearson test (if n>50) or Shapiro-Wilk test (if n<50). When the data followed a normal distribution, we then compared them using parametric Student t-tests (between 2 groups) or one One-way ANOVA (between >2 groups). When data did not follow a normal distribution, we compared them using Mann-Whitney (2 groups) or Kruskal-Wallis (>2 groups) non-parametric tests. p = 0.0001 - 0.001 or <0.0001 was considered extremely significant (***), p = 0.001 - 0.01 was considered very significant (**); p = 0.01 - 0.05 was considered significant (*) and p = ≥0.05 was considered not significant (ns). Number of embryos and experiments performed for all analysis are listed in table 5.

**Table 5:** Experimental replicates and sample sizes.

| Experiment | Condition | Number of MCCs<br>(or BBs for TEM)<br>analyzed | Number of<br>embryos<br>analyzed | Number of<br>experimental<br>repeats |
| --- | --- | --- | --- | --- |
| **BB orientation** | Control | 125 | 4-9 | 2-3 |
|  | Lrrcc1 ATG-MO | 24 |  |  |
|  | Lrrcc1 Spl-MO | 28 |  |  |
|  | Ccdc61 ATG-MO | 33 |  |  |
|  | Ccdc61 Spl-MO | 27 |  |  |
|  | Lrrcc1 ATG-MO +mRNA | 40 |  |  |
|  | Lrrcc1 Spl-MO +mRNA | 45 |  |  |
|  | Ccdc61 ATG-MO +mRNA | 39 |  |  |
|  | Ccdc61 Spl-MO +mRNA | 20 |  |  |
| **MCC**<br>**polarization**<br>**Rayleigh Test** | Control | 591 | 4-9 | 3 |
|  | Lrrcc1 ATG-MO | 181 |  |  |
|  | Lrrcc1 Spl-MO | 33 |  |  |
|  | Ccdc61 ATG-MO | 158 |  |  |
|  | Ccdc61 Spl-MO | 159 |  |  |
| **BB docking** | Control | 28 | 3-9 | 2-3 |
|  | Lrrcc1 ATG-MO | 36 |  |  |
|  | Lrrcc1 Spl-MO | 33 |  |  |
|  | Ccdc61 ATG-MO | 34 |  |  |
|  | Ccdc61 Spl-MO | 28 |  |  |
|  | Lrrcc1 ATG-MO +mRNA | 41 |  |  |
|  | Lrrcc1 Spl-MO +mRNA | 35 |  |  |
|  | Ccdc61 ATG-MO +mRNA | 35 |  |  |
|  | Ccdc61 Spl-MO +mRNA | 29 |  |  |
| **BB spacing** | Control | 19 | 2 | 2-3 |
|  | Lrrcc1 ATG-MO | 20 |  |  |
|  | Lrrcc1 Spl-MO | 28 |  |  |
|  | Ccdc61 ATG-MO | 27 |  |  |
|  | Ccdc61 Spl-MO | 11 |  |  |
|  | Lrrcc1 ATG-MO +mRNA | 43 |  |  |
|  | Lrrcc1 Spl-MO +mRNA | 27 |  |  |
|  | Ccdc61 ATG-MO +mRNA | 28 |  |  |
|  | Ccdc61 Spl-MO +mRNA | 28 |  |  |
| **Actin signal** | Control (of Lrrcc1 ATG) | 72 | 3 | 2 |
|  | Control (of Lrrcc1 Spl) | 43 |  |  |
|  | Lrrcc1 ATG-MO | 31 |  |  |
|  | Ccdc61 ATG-MO | 32 |  |  |
| **Flow engagement** | Control |  | 8-12 | 1-2 |
|  | Lrrcc1 ATG-MO |  |  |  |
|  | Ccdc61 ATG-MO |  |  |  |
| **Cilia beating** | Control | 16 | 6-9 | 1 |
|  | Lrrcc1 ATG-MO | 23 |  |  |
|  | Ccdc61 ATG-MO | 13 |  |  |
| **Pericentrin signal** | Control (of Lrrcc1) | 91 | 5-9 | 2-3 |
|  | Lrrcc1 ATG-MO | 136 |  |  |
|  | Control (of Ccdc61) | 46 |  |  |
|  | Ccdc61 ATG-MO | 143 |  |  |
| **Rootlet presence (TEM)** | Control | 134 | 2-3 | 1-2 |
|  | Lrrcc1 ATG | 174 |  |  |
|  | Ccdc61 ATG | 120 |  |  |
| **Embryo survival in presence of bacteria** | Control |  | 73 | 3 |
|  | Control+BACT |  | 84 |  |
|  | GFP |  | 73 |  |
|  | GFP +BACT |  | 74 |  |
|  | Lrrcc1 ATG |  | 45 |  |
|  | Lrrcc1 ATG +BACT |  | 53 |  |
|  | Ccdc61 ATG |  | 35 |  |
|  | Ccdc61 ATG +BACT |  | 38 |  |

